# Antibodies Expand the Scope of Angiotensin Receptor Pharmacology

**DOI:** 10.1101/2023.08.23.554128

**Authors:** Meredith A. Skiba, Sarah M. Sterling, Shaun Rawson, Morgan S.A. Gilman, Huixin Xu, Genevieve R. Nemeth, Joseph D. Hurley, Pengxiang Shen, Dean P. Staus, Jihee Kim, Conor McMahon, Maria K. Lehtinen, Laura M. Wingler, Andrew C. Kruse

## Abstract

G protein-coupled receptors (GPCRs) are key regulators of human physiology and are the targets of many small molecule research compounds and therapeutic drugs. While most of these ligands bind to their target GPCR with high affinity, selectivity is often limited at the receptor, tissue, and cellular level. Antibodies have the potential to address these limitations but their properties as GPCR ligands remain poorly characterized. Here, using protein engineering, pharmacological assays, and structural studies, we develop maternally selective heavy chain-only antibody (“nanobody”) antagonists against the angiotensin II type I receptor (AT1R) and uncover the unusual molecular basis of their receptor antagonism. We further show that our nanobodies can simultaneously bind to AT1R with specific small-molecule antagonists and demonstrate that ligand selectivity can be readily tuned. Our work illustrates that antibody fragments can exhibit rich and *evolvable* pharmacology, attesting to their potential as next-generation GPCR modulators.

## Introduction

G protein-coupled receptors (GPCRs) are central regulators of nearly all aspects of human physiology. Decades of pharmacological investigation and hundreds of structural studies have illustrated how chemically distinct ligands stimulate or suppress G protein-coupled receptor signaling. Collectively GPCRs remain one of the most successful classes of therapeutic drug targets, with four new GPCR-targeted pharmaceuticals approved in the past year alone^1,2^. However, the lack of subtype and tissue selectivity of GPCR-targeted small-molecule and peptide ligands remains a challenge for therapeutically relevant receptors, both in the clinic and in research settings.

Antibodies provide a potential solution to these challenges. In contrast to conventional ligands, antibodies are expected to interact with GPCRs in fundamentally different ways and to exhibit increased subtype selectivity through interactions with extended epitopes outside of the endogenous ligand-binding orthosteric pocket^3^. High specificity combined with antibody engineering strategies could provide improved approaches to drugging even well-established therapeutic targets. For example, undesired side effects could be reduced with bispecific antibodies that modulate GPCR signaling only in specific cell types, pharmacokinetics can be tuned to control circulating half-life through antibody constant region (Fc) engineering, and antibody effector functions can be evoked to target cells for ablation by promoting immune-mediated cell death^4^. In addition, GPCR-targeting antibodies can be incorporated into chimeric antigen receptor (CAR) T-cell technologies to deplete cells expressing GPCRs^5^, or may be integrated with targeted protein degradation technologies to inactivate otherwise undruggable GPCRs. In the laboratory, antibodies can be expressed in single cell types in model organisms or within intracellular organelles to provide tools to interrogate the role of GPCR signaling in various physiological processes, as illustrated by antibody fragment biosensors^6–8^.

Despite their promise, the properties of antibodies as GPCR ligands have not yet been characterized in-depth. We therefore sought to assess whether antibodies can modulate the angiotensin II type I receptor (AT1R), a prototypical family A GPCR, in fundamentally new ways to address therapeutic gaps in maternal health left by small molecule drugs. AT1R modulates renal and cardiovascular function in response to the eight amino acid peptide hormone angiotensin II (AngII). AT1R is one of the most successfully drugged GPCRs by small molecules, with small-molecule angiotensin receptor blockers (ARBs) being commonly prescribed to reduce high blood pressure^9^. However, these pharmacological interventions cannot be used to treat renal and hypertensive disorders during pregnancy such as preeclampsia, as the small molecules readily cross the placenta and cause on-target fetotoxicity^10^. Other frontline anti-hypertensives, including angiotensin-converting enzyme (ACE) inhibitors, are similarly fetotoxic and contraindicated during pregnancy, leaving few options for pregnant patients. Proteins are too large to cross the placental barrier and could therefore provide an avenue to safely modulate the maternal renin-angiotensin system during pregnancy^11^.

Previously, we described a series of heavy chain-only antibody (nanobody) antagonists for the angiotensin II type I receptor (AT1R) that we isolated from a synthetic library^12,13^. Here, we show that it is possible to evolve and engineer nanobody antagonists specific for targeting maternal AT1R and demonstrate that they can be restricted to maternal circulation *in vivo*. We then use structural approaches to examine how nanobodies engage with the extracellular face of AT1R to modulate signaling, which is controlled by allosteric networks deep within the receptor’s transmembrane core. We found that nanobodies that are closely related in sequence can have profoundly divergent pharmacological properties. Collectively, our data demonstrate that antibody scaffolds have rich pharmacological capacity and are well-suited to act as both competitive and allosteric modulators of GPCRs.

## Results

### Evolving and engineering a nanobody antagonist to target maternal AT1R

Previously, we discovered the AT118 family of nanobody antagonists that target AT1R. The parent nanobody, AT118, binds competitively with both the peptide agonist AngII and the small-molecule antagonist olmesartan^13^. These competitive interactions are also seen with AT118-A, a high-affinity variant of AT118, and AT118-H, a humanized high-affinity variant (Fig. 1a, Supplementary Table 1). Functionally, AT118-A suppresses AT1R signaling in cellular assays and reduces blood pressure in mice, with similar potency to clinically approved antagonists^13^. An AT118 derivative could in principle be employed to selectively target maternal AT1R during pregnancy, as transport across the placenta is largely governed by size, but current derivatives require further improvement for viable therapeutic applications. For example, at a molecular weight of 15 kDa, a nanobody alone would be cleared via renal filtration in minutes, thereby limiting therapeutic potential^14^. In addition, AT118-H displays high levels of off-target binding, which correlates with a short circulating half-life^15,16^.

**Figure 1.**
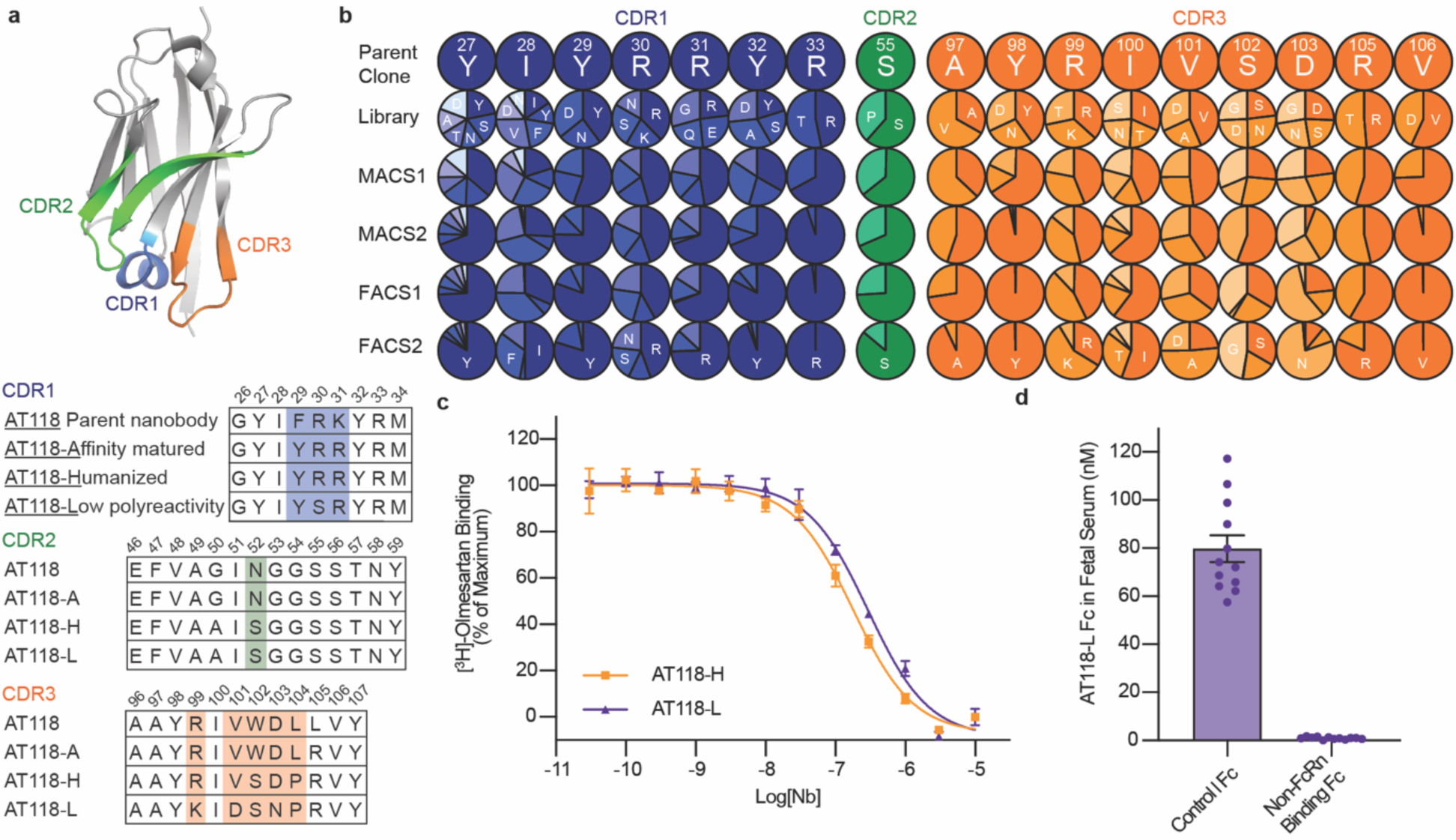
Evolution of AT118 family of nanobody antagonists. a) Structure of AT118-H (PDB 7T83)^16^ and CDR sequences of nanobodies described in this study colored by CDR. Shaded positions vary between nanobodies. Full nanobody sequences are provided in Supplementary Table 1. b) Sites substituted in AT118-H library. Variant enrichment was tracked through each selection round via deep sequencing. c) AT118-H and evolved variant AT118-L displace olmesartan from cell membranes containing AT1R. Error bars represent mean ± standard error from three experiments. d) AT118-L Fc fusion proteins lacking the ability to interact with FcRn (pMAS493, Supplementary Table 1) do not accumulate in fetal serum of mice compared to AT118-L Fc fusion proteins with a WT Fc (pMAS430, Supplementary Table 1). Error bars represent mean ± standard error from twelve embryos from three separate litters.

We first sought to evolve a variant of AT118-H with increased specificity for AT1R. We mutagenized seventeen positions of AT118-H complementarity-determining regions (CDRs) to create a yeast-display library of 5×10^9^ AT118-H variants (Fig. 1b). After bulk enrichment of AT1R binders by magnetic-activated cell sorting (MACS), we conducted a FACS selection to isolate clones that bound to AT1R with high specificity. We further enriched for high-affinity AT1R binding clones in a final selection round (Supplementary Data Figs. 1a-e). In the affinity driven selection, CDR3 was more amenable to mutagenesis than CDR1 (Fig. 1b). We combined tolerated mutations to generate a panel of AT118 variants that were screened for on-target binding to AT1R and non-specific binding on yeast (Supplementary Data Fig. 1f-g). All variants maintained strong binding to detergent-solubilized AT1R. Introduction of R30S^CDR1^ and V101D^CDR3^ substitutions yielded variants with reduced non-specific binding [superscripts refer to nanobody CDR] (Supplementary Data Fig. 1g)^16^.

We then evaluated pharmacological function by testing if the evolved variants displace the small-molecule antagonist olmesartan from AT1R’s orthosteric pocket like the parent nanobody AT118-H (Supplementary Data Fig. 2a-c). S102G^CDR3^ and D103N^CDR3^ variants bound strongly to AT1R but lost the ability to displace olmesartan (Supplementary Data Fig. 2a). However, the ability to compete with olmesartan binding was rescued by combining the V101D^CDR3^ substitution with D103N^CDR3^ (Supplementary Data Fig. 2b). We concluded that pharmacological function is largely influenced by CDR3, whereas CDR1 and 2 play a stronger role in receptor recognition. Ultimately, we selected the AT118-H R30S, R99K, V101D, D103N variant (AT118-L), which exhibited low non-specific binding but retained the affinity and pharmacological function of AT118-H, for further experiments (Fig. 1c, Supplementary Data Fig. 2d).

We further tuned the pharmacokinetics of AT118-L to selectively target maternal AT1R through antibody engineering. To extend circulating half-life, we fused AT118 to an IgG1 Fc, dimerizing the nanobody and raising the molecule’s molecular weight above the ∼70 kDa renal filtration cutoff^17^. To retain AT118-L in maternal circulation, we mutated the Fc’s neonatal Fc receptor (FcRn) binding site, eliminating the active transport mechanism for antibodies across the placenta^18,19^. Additionally, we blocked undesired cytotoxic antibody effector functions by inhibiting Fc gamma receptor binding and complement fixation through established mutations^20^. The AT118-L Fc fusion protein retains the ability to bind AT1R and acts as an antagonist in cellular signaling assays (Supplementary Data Fig. 2d, Supplementary Table 2).

To test if AT118-L is selectively targeted to maternal tissues, we treated pregnant mice with the fusion proteins and analyzed levels of AT118-L Fc in fetal serum. Consistent with specificity in targeting, our engineered AT118-L Fc fusion proteins were minimally detected in fetal serum, whereas control Fc fusions that support FcRn binding and transport across the placenta accumulated in fetal serum (Fig. 1d). An alternative AT118 variant, lacking the capacity to bind to AT1R, displays similar fetal serum levels (Supplementary Data Fig. 2e).

### Determining nanobody-receptor complex structure

The discrepancy between AT1R binding and ability to displace olmesartan from AT1R observed in the evolved AT118-H variants was puzzling, raising the possibility that the AT118 nanobody family may have unexpected properties. Furthermore, prior double electron-electron resonance (DEER) spectroscopy studies indicate that AT118-A and small-molecule antagonists stabilize distinct inactive states of AT1R, suggesting that nanobodies antagonize AT1R through a unique mechanism^13^. To gain insight into the mechanism of action of the AT118 series of nanobodies, we sought to determine the structure of AT118-H fused to AT1R. Because crystallographic approaches using the lipidic cubic phase (LCP) method did not generate crystals that diffracted to sufficient resolution (Supplementary Data Fig. 3) and the 55 kDa AT118-H AT1R complex failed to produce high-quality cryo-electron microscopy (cryo-EM) reconstructions, we collected small cryo-EM data sets of our nanobody AT1R complex with a series of adaptors to increase mass and aid in particle alignment (Supplementary Data Fig. 3, Supplementary Table 3)^21–24^. We found that an AT1R thermostabilized apocytochrome b562 (BRIL) fusion protein complexed with an anti-BRIL Fab and anti-Fab nanobody was the most successful approach to obtain high-quality structural data. We determined a structure of the AT118-H-AT1R-BRIL fusion protein in complex with anti-BRIL Fab and anti-Fab nanobody to a global resolution of 2.9 Å (Fig. 2a, Supplementary Data Fig. 4, Supplementary Table 4). Density was well defined for both AT1R and AT118-H and local refinement yielded resolutions of 2.7 Å at the AT118 AT1R interface (Supplementary Data Fig. 4). AT118-H engages with the extracellular region of AT1R, with all three CDR regions interacting with AT1R’s extracellular loops (ECLs) (Fig. 2b).

**Fig 2.**
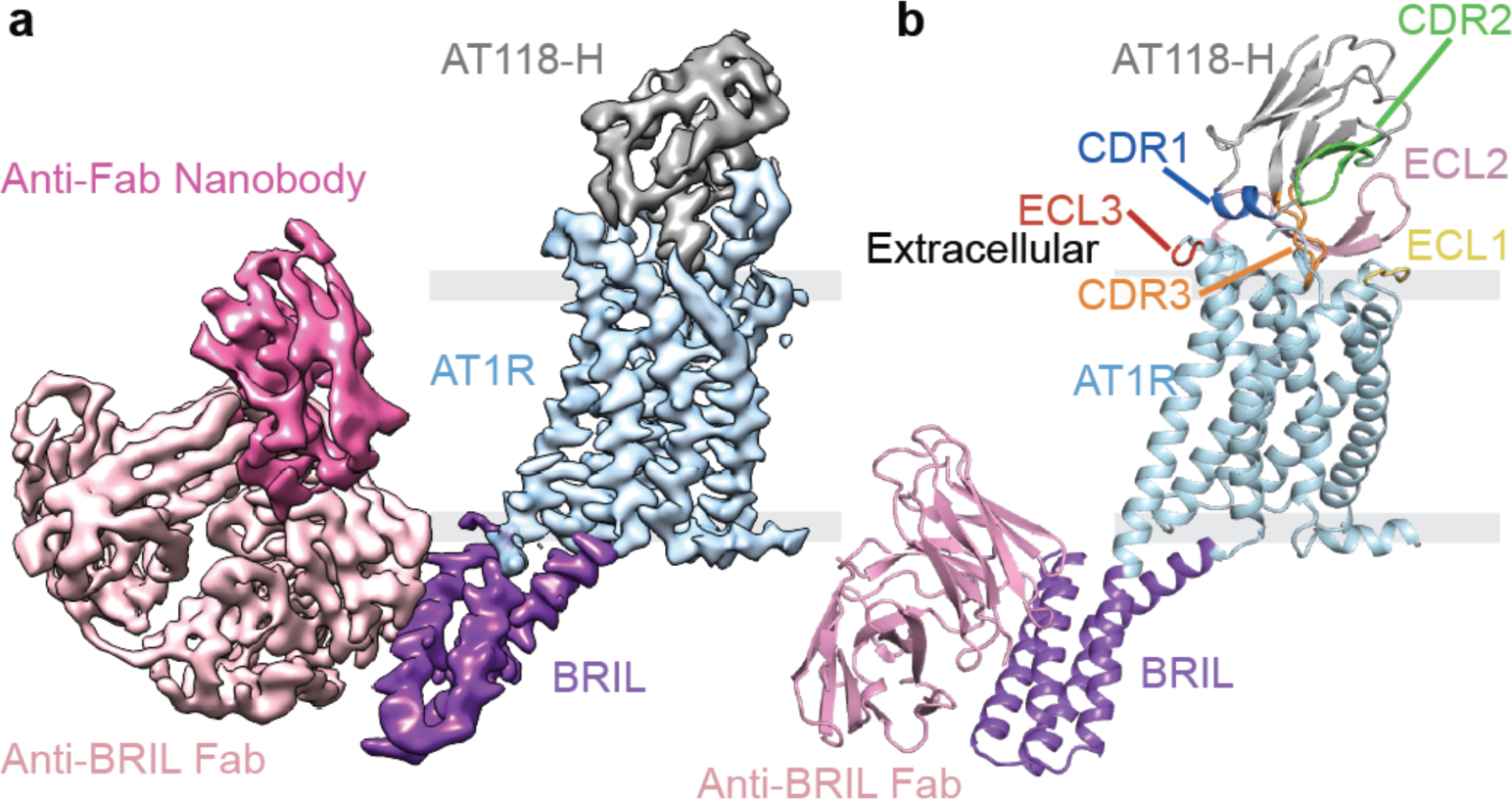
Structure of H bound to AT1R. a) Cryo-EM map colored by component: AT118-H (gray), AT1R (light blue), BRIL (purple), anti-BRIL Fab (light pink), anti-Fab nanobody (dark pink) b) Model of the AT118-H AT1R complex from local refinement maps. Image is colored as in a) with extracellular loops and AT118-H CDRs colored as follows: ECL1 (yellow), ECL2 (light pink), ECL3 (red), CDR1 (dark blue), CDR2 (green), CDR3 (orange).

### Antagonist AT118-H mimics many of the interactions formed by peptide agonists

Despite acting as an antagonist, AT118-H shares a similar binding mode with peptide agonists and stabilizes the extracellular face of the receptor in an active-like conformation. CDR3 occupies AT1R’s orthosteric pocket and mimics the binding of a peptide ligand, forming β-strand interactions with ECL2 (Fig. 3a-f)^25–27^. AT118-H I100^CDR3^ resemble peptide ligand AngII residue V3 and participates in hydrophobic interactions with I172^ECL2^ and V179 ^ECL2^ (Fig. 3d, 3e) [superscripts indicate extracellular loop locations in AT1R and Ballesteros Weinstein numbering for conserved GPCR residues^28^]. This hydrophobic network is further supported by AT1R residues Y87^2.63^, Y92^ECL1^, and F170^ECL2^ and Y98^CDR3^ in AT118-H (Fig. 3d). Maintaining these peptide-mimicking hydrophobic interactions is critical for AT118-H binding, as disruption of this network with smaller or polar residues is largely not tolerated in our AT118-H protein engineering campaign or direct binding assays (Fig. 1b, 3g). Additional residues in ECL1 are not critical for AT118-H binding, as ECL1 can be replaced with the equivalent ECL from the related angiotensin II type II receptor (AT2R) (Fig. 3g).

**Fig 3.**
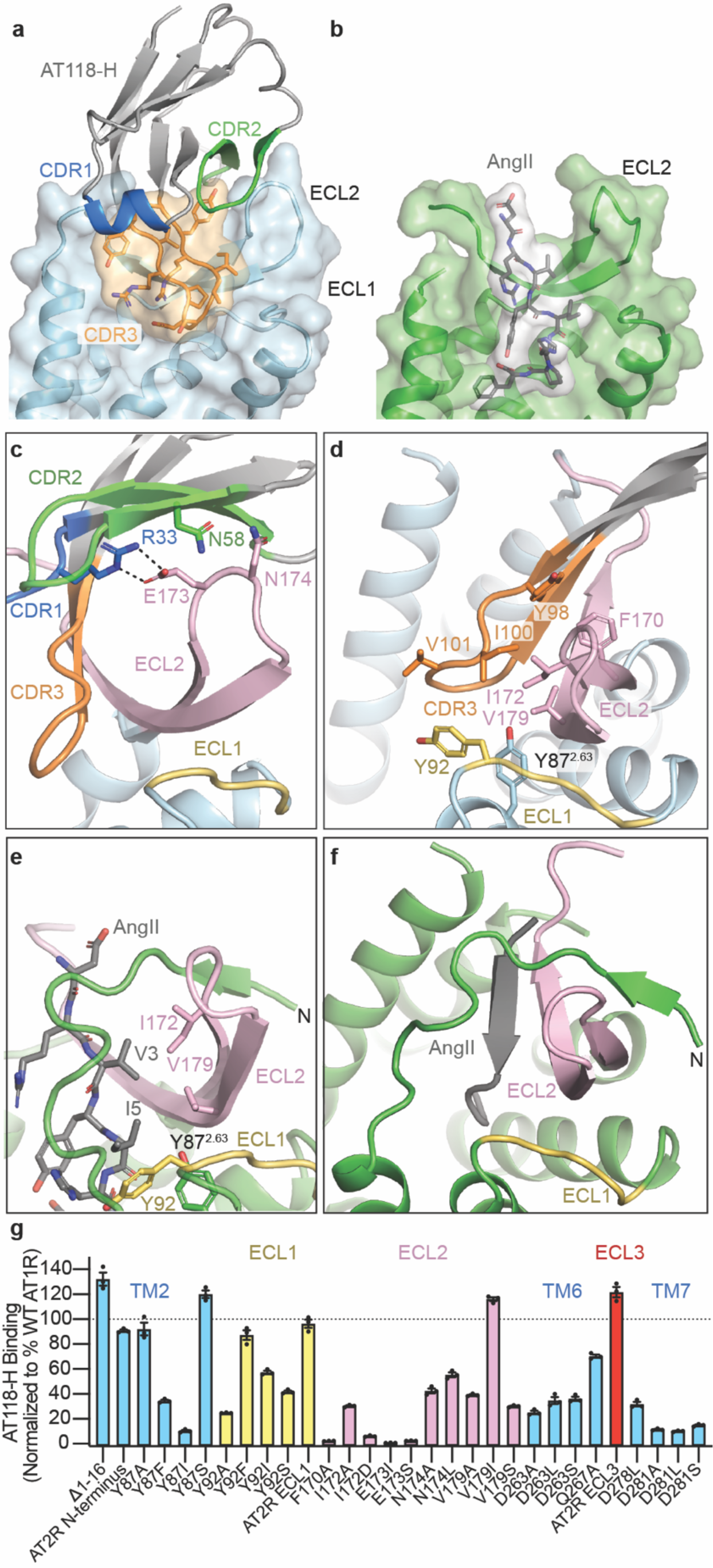
AT118-H mimics peptide agonist binding to AT1R. a) Surface rendering of AT118-H bound to AT1R as colored in Fig 2b. CDR3 (orange) engages with the orthosteric pocket of AT1R. b) surface rendering of active-state AT1R bound to the endogenous peptide agonist AngII (gray) (PDB 6OS0)^26^. AngII engages deeply in the orthosteric pocket. c) AT118-H binding is driven by engagement of CDRs 2 (green) and 3 (orange) with ECL2 (pink). Dashed lines indicate hydrogen bonds. d) AT118-H CDR3 (orange) forms β-strand and hydrophobic interactions with ECL2 (pink). The hydrophobic network is further supported by residues on ECL1 and TM2. e) AngII (gray sticks) and CDR3 share similar hydrophobic interactions with ECL2 (pink). f) In peptide agonist bound structures, the N-terminus of AT1R stabilizes the position of ECL2 through β-strand interactions. g) Binding of AT118-H to Expi293 cells expressing AT1R variants. Error bars represent mean ± standard error from three experiments. In the AT2R N-terminus chimera residues 2-16 of AT1R are replaced with residues of 3-33 of human AT2R. The AT2R ECL1 chimera contains M90Y, E91R, R93D, and P95L substitution and ECL3 chimera contains I270V, R272N, D273S substitutions.

The binding mode of AT118-H displaces the N-terminus of AT1R from the extracellular face of the receptor, leaving it disordered. In active-state structures, the N-terminus of AT1R wraps around AT1R’s ECL2 β-hairpin and makes contacts with both the N-terminal β-strand and β-turn stabilizing the peptide agonist binding site (Fig. 3e-f)^25–27^. While the unraveling of this structural feature by AT118-H is surprising, AT1R’s N-terminus is not critical for AT118-H binding, as the nanobody binds strongly to AT1R when the N-terminus is truncated or replaced (Fig. 3g). Instead, CDR2 imitates the receptor’s N-terminus and stabilizes the position of ECL2 (Fig. 3c, 3e). R33^CDR1^ anchors AT118-H onto the β-turn of ECL2 through interactions with E173^ECL2^ (Fig. 3c). This interaction, together with a neighboring interaction between N174^ECL2^ and the backbone of CDR2 and N58^CDR2^, are critical for nanobody binding (Fig. 1b, Fig. 3g).

AT118-H mimics agonist binding beyond the peptide binding site, inducing outward movement of ECL3 and the extracellular side of transmembrane helices (TMs) 6 and 7 and corresponding inward movement of TM5 (Fig. 4a-d). This movement repositions K199^5.42^ and constricts AT1R’s ligand binding pocket, as observed in active-state structures (Fig. 4d). ECL3 is not essential for stabilizing the TM 5/6/7 rearrangement, as an AT1R-AT2R ECL3 chimera fully accommodates AT118-H binding (Fig. 3g). Instead, a compatible binding interface between positively charged residues in AT118H’s CDRs and the adjacent Asp rich TMs drives this movement. R99^CDR3^ directly interacts with D281^7.32^ and mimics a key interaction between R2 of peptide agonists and TM7 required for full agonism by AngII (Fig. 4a-b)^25–27,29^. This interaction is required for AT118-H binding, as substitution of R99^CDR3^ and nearby R31^CDR1^ with residues other than lysine and replacement of D263^6.58^, D278^7.29^, and D281^7.32^ suppress binding of the nanobody to AT1R (Fig. 1b, 3g). Overall, this larger scale conformational change may correspond to similarities in DEER spectroscopy measurements of AT118-A and TRV026, a β-arrestin biased peptide agonist, bound AT1R^13^. Like AT118-A, TRV026 induces some but not all of the conformational changes in AT1R that are required for full activation of the receptor^26,27^.

**Fig 4.**
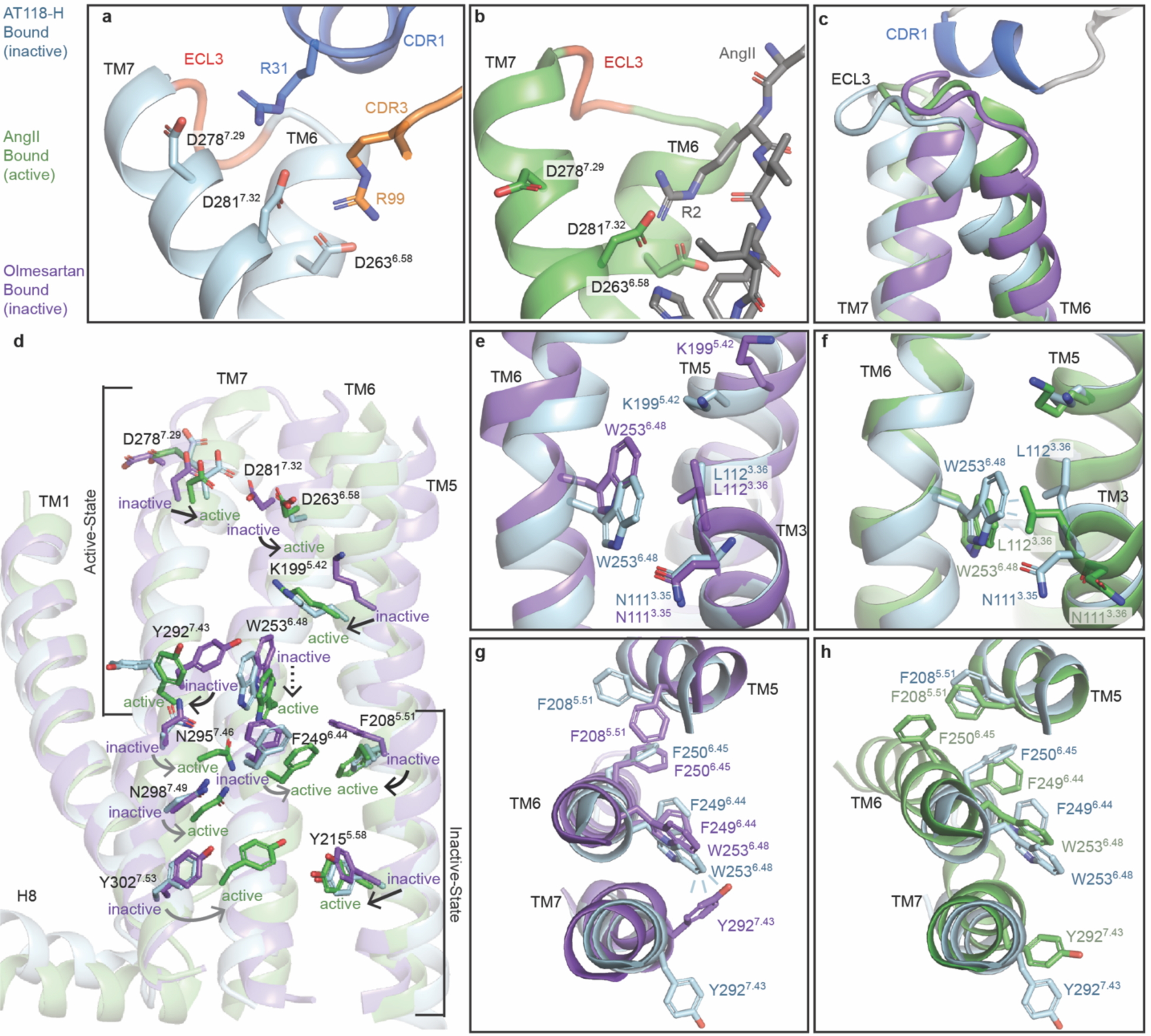
AT118-H stabilizes a hybrid active-inactive state of AT1R. a-b) R31^CDR1^ (a, blue sticks) and R99^CDR3^ (a, orange sticks) of AT118-H mimic R2 of AngII (b, gray sticks) positioning the extracellular side of Asp rich TMs 6 and 7. c) TM6, ECL3, and TM7 are displaced outward in AT118-H (light blue) and AngII (green) bound AT1R compared to small-molecule antagonist bound AT1R (purple) (PDB 4ZUD)^30^. d) AT118-H partially engages the allosteric activation network between AT1R’s orthosteric pocket and the intracellular side of AT1R. Switches indicated with the following arrows are activated (black), partially activated (dashed lines), unengaged (gray). e) AT118-H induced movements of TM5 and TM6 (light blue) induces inward movement of K119^5.42^ and downward rotation of W253^6.48^ compared to the inactive state (purple). f) W253^6.48^ does not fully rotate into its active-state position (green) and cannot displace N111^3.35^ and L112^3.36^ g) Partial movement of W253^6.48^ repositions Y292^7.43^ and F208^5.51^ from the canonical inactive state (purple) to h) active state positions (green), but is not sufficient to rotate the F249^6.44^ and F250^6.45^ rachet and displace TM6 to open up the intracellular effector binding site.

### AT118-H retains AT1R in an inactive state

AT118-H induces many conformational changes that are also present in active-state AT1R structures, but acts as an antagonist because it fails to propagate conformational change through the receptor core to activate the intracellular side of the receptor (Fig. 4d). AT118-H CDR1 displaces the extracellular side of TM6 further than peptide agonists and restricts rotation of the helix to its fully activated position (Fig. 4a, 4c, 4d). This partial rotation facilitates downward movement of W253^6.48^ to an intermediate position between the inactive and active state (Fig. 4d, 4e, 4f). Partial movement of W253^6.48^ causes a steric clash with the Y292^7.43^ side chain, inducing rotation of Y292^7.43^, as seen in structures of AT1R bound to the endogenous agonist AngII and high affinity Sar1-AngII derivative (Fig. 4g)^26,27^. While movement of Y292^7.43^ is a critical step required for G-protein binding, AT118-H does not induce all of the movements required for outward displacement of TM6 to accommodate transducer binding. This includes rotation of L81^2.57^ in the base of the orthosteric pocket, F249^6.44^ and F250^6.45^ in the phenylalanine rachet, N111^3.35^, N295^7.46^ and N298^7.49^ in the canonical sodium binding site, and Y302^7.53^ at the water lock (Fig. 4d-h)^25,26^. This intermediate conformational state of AT1R is driven by the higher binding position of AT118-H in the orthosteric pocket as compared to peptide agonists. This binding mode fails to fully engage the allosteric network within the transmembrane core to transition AT1R to the active state.

### AT118-H derivatives display probe dependence for small-molecule antagonists

The shallow binding mode of AT118-H leaves the small-molecule antagonist binding site deep within the orthosteric pocket vacant, with only weak density observed for the side chain of W84^2.60^ beneath CDR3 (Fig. 3a, Supplementary Data Fig. 5a, 5b). Comparison with inactive-state structures suggested that some small-molecule antagonists could be accommodated below CDR3, whereas others like olmesartan may clash with CDR3 and compete with AT118-H binding (Fig. 1c, Supplementary Data Fig. 5b)^30,31^. We examined binding of AT118 family members to AT1R with a broad series of small-molecule antagonists (Supplementary Data Fig. 5c). Nearly all small-molecule antagonists of AT1R contain a core biphenyl, with one side conjugated to a tetrazole or oxadiazole, and the other side decorated with a diverse variety of functionalities (Supplementary Data Fig. 5a). We found that AT118-H and AT118-L fully compete for binding to AT1R with peptide ligands but exhibit tolerance of small-molecule antagonist binding to varying degrees (Supplementary Data Fig. 5c). Small antagonists bound simultaneously with AT118-H and AT118-L, whereas antagonists with large bulky substituents were the least compatible with nanobody binding due to predicted steric clashes with CDR3 (Supplementary Data Fig. 5a, 5d). This selectivity, referred to as “probe dependence”, is commonly observed for small-molecule allosteric modulators^32,33^. Intriguingly, both AT118 variants fully tolerate binding of losartan, but not olmesartan, which is closely related in chemical structure (Supplementary Data Fig. 5a). We hypothesized that changes in nanobody probe dependence may explain the discrepancy between AT1R binding and the ability to displace olmesartan that we encountered in our protein engineering campaign (Supplementary Data Fig. 2a-c).

### CDR3 tunes small molecule probe dependence

To investigate the relationship between probe dependence and CDR3 sequence, we determined a structure of AT118-L and losartan in complex with the AT1R-BRIL fusion protein, anti-BRIL Fab, and anti-Fab nanobody to a global resolution of 3.3 Å (Fig 5a, Supplementary Data Fig. 6). Density for losartan within the orthosteric pocket is unambiguous. AT118-L and AT118-H exhibit an identical binding mode and stabilize AT1R is a similar conformation (0.67 r.m.s.d. over 237 AT1R C⍺ atoms). Overall, AT118-L’s CDR3 engages more deeply with the orthosteric pocket than AT118-H (Fig. 5a). The depth of CDR3 binding is largely driven by interactions with TM7 (Fig. 5a-c). In AT118-H, the side chain of S102^CDR3^ hydrogen bonds with the backbone of D281^7.32^, which restrains CDR3 within the orthosteric pocket (Fig. 5b). The V101D^CDR3^ substitution found in AT118-L forms an analogous hydrogen bond with the D281^7.32^ backbone carbonyl (Fig. 1a, Fig. 5c-d).

**Fig 5.**
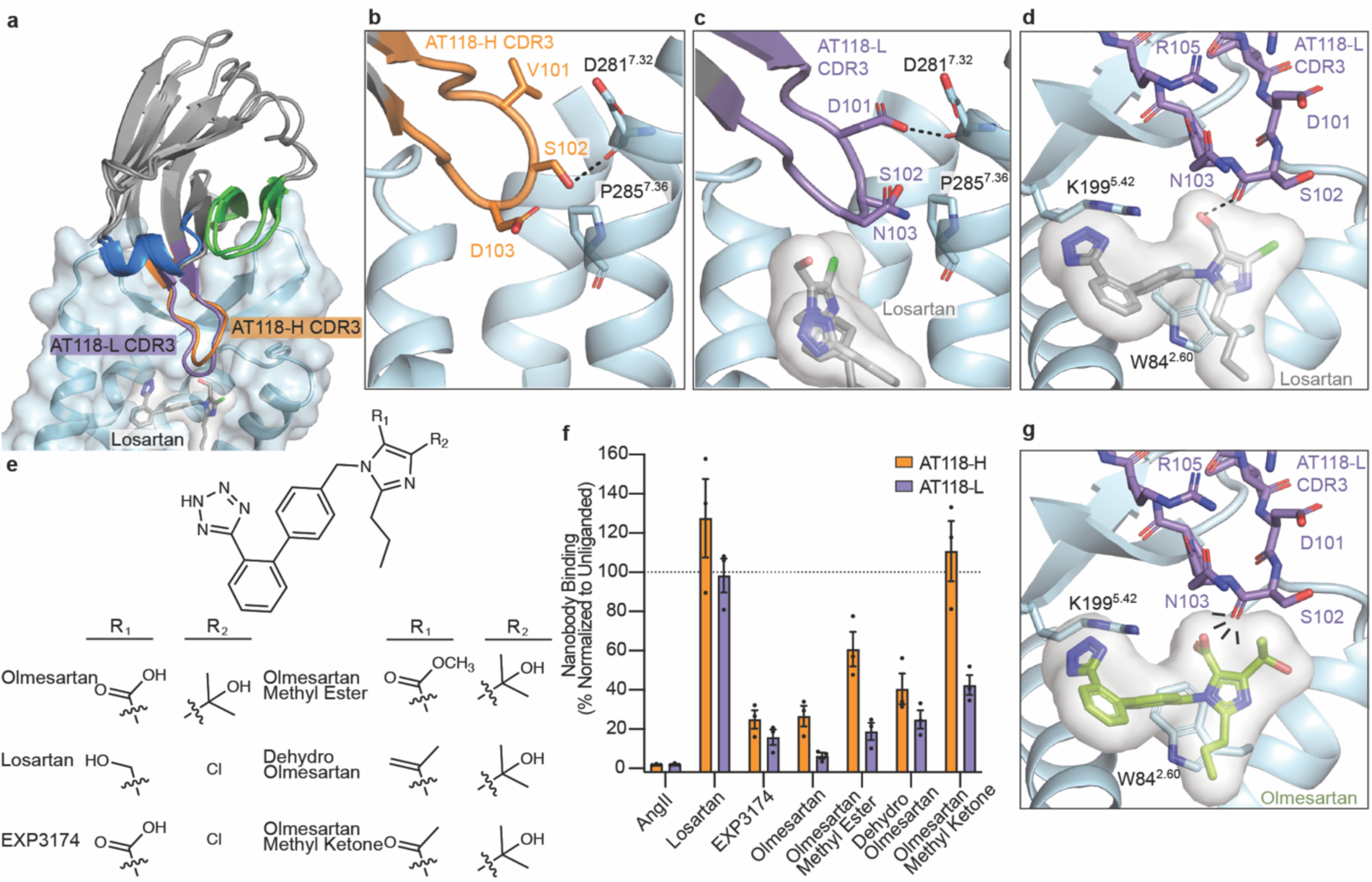
AT118 family members exhibit probe dependence with small molecules. a) AT118-H and the evolved AT118-L variant accommodate binding of the small-molecule antagonist losartan (gray sticks). AT118-L CDR3 (purple) engages deeper within the orthosteric pocket compared to AT118-H (orange). b) S102^CDR3^ of AT118-H and c) D101^CDR3^ of AT118-L controls the depth of CDR3 within the orthosteric pocket through interactions with TM7. d) The backbone carbonyl of S102^CDR3^ of AT118-L is within hydrogen binding distance of the alcohol substituent (R_1_ in e) of losartan (gray sticks). e) Olmesartan and losartan derivatives vary in substituents on the imidazole ring. f) Probe dependent binding of AT118-H (orange) and AT118-L (purple) with losartan and olmesartan derivatives. Error bars represent mean ± standard error from three experiments. g) CDR3 of AT118-L clashes with the carboxylic acid (R_1_ in e) of modeled olmesartan (green sticks, modeled based upon PDB: 4ZUD).

We then examined the molecular basis for the divergent binding properties of losartan and olmesartan, which share the same core scaffold and differ only in substituents on the imidazole (Fig. 5e, Supplementary Data Fig. 5a). In our structure, the backbone carbonyl of AT118-L S102^CDR3^ hydrogen bonds with losartan’s alcohol substituent (Fig. 5d). Replacement of the alcohol with the more acidic carboxylic acid found in the equivalent position of olmesartan eliminates the favorable hydrogen bond. Instead, a repulsive delocalized negative charge is introduced, displacing olmesartan from the orthosteric pocket. D103^CDR3^ of AT118-H would also clash with olmesartan’s carboxylic acid, accounting for AT118-H’s ability to compete with olmesartan (Supplementary Data Fig. 5b). This observation is further supported by the loss of pharmacological activity associated with the D103N^CDR3^ substitution in our protein engineering campaign (Supplementary Data Fig. 2a-c). AT118-H D103N^CDR3^ variants tolerate olmesartan binding, as the clash between D103^CDR3^ and olmesartan’s carboxylic acid is eliminated (Supplementary Data Fig. 5b). However, the introduction of V101D^CDR3^ leads to repositioning of CDR3, as observed in our AT118-L structure, yielding a CDR3 binding mode that is incompatible with olmesartan binding, as demonstrated by the competitive behavior of AT118-H V101D^CDR3^ D103N^CDR3^ containing variants (Fig. 5f-g, Supplementary Data Fig. 2b, 2c).

To directly compare the influence of alcohol and carboxylic acid substituents on probe dependence, we tested binding of AT118-H and AT118-L to AT1R in the presence of EXP3174, an active metabolite of losartan in which the alcohol is converted to a carboxylic acid (Fig. 5e-f). Like olmesartan, EXP3174 inhibits AT118-H and AT118-L binding to AT1R (Fig. 5f). To further probe the role of the carboxylic acid, we evaluated a series of olmesartan derivatives, where the carboxylic acid is replaced with less acidic substituents (Fig. 5e). Conversion of the carboxylic acid to a methyl ester or an alkene modestly increased AT118-L and AT118-H binding, whereas replacement with methylketone increased binding four-fold (Fig. 5f). In the case of AT118-H, the methylketone derivative fully supports nanobody binding, which is likely due to the shallower binding mode of AT118-H relative to AT118-L. Overall, our work demonstrates that the AT118 family of nanobodies are competitive antagonists that prevent binding of peptide ligands but can also function as probe-dependent negative allosteric modulators that support binding of some small-molecule AT1R antagonists.

## Discussion

Most therapeutic antibodies function by blocking protein-ligand interactions or directing immune effectors to a protein of interest. However, few antibodies are reported to functionally interact with GPCRs. We demonstrated that antibody fragments can interact with GPCRs in unusual ways to directly alter receptor signaling and exert pharmacological effects on multiple levels that are not readily achievable by small molecules. Our work illustrates that antibodies are poised not just to achieve enhanced receptor and tissue selectivity, but to also exert distinct allosteric effects and stabilize unique receptor conformations that influence signaling outputs.

We probed the mechanistic basis for antibody-mediated antagonism of AT1R, a prototypical GPCR. All members of the AT118 series of nanobodies engage with AT1R’s ECL2 and directly compete for binding with peptide agonists (Fig. 3c, 3d, 5a, 5f). This binding mode differs from functional antibodies described for chemokine, protease-activated (PAR) receptors and family B GPCRs, where antibodies modulate signaling through interactions with extended N-termini and large ectodomains^34–43^. Interestingly, ECL2 is the reported epitope for naturally occurring self-reactive AT1R-targeting autoantibodies associated with preeclampsia, suggesting that it is an immunogenic hot-spot^44,45^.

Strikingly, AT118-H and AT118-L mimic peptide agonist binding and stabilize the extracellular face of AT1R in an active conformation but do not engage with key activation switches deep within the orthosteric pocket of AT1R, leaving the intracellular region in an inactive state (Fig. 4d). Stabilization of this unusual hybrid conformation suggests that binding of antibodies to extracellular loops can trap distinct states of GPCRs that differ from conventional ligands that bind within the orthosteric pocket, which may influence signaling output in distinct ways. For example, recent studies suggest that G-protein selectivity can be influenced by ligand interactions with extracellular loops^46^. Overall, these findings highlight that ligands can employ diverse molecular mechanisms to achieve similar functional effects. This has implications for rational drug design and could broaden the conformational space explored during such efforts.

By simultaneously interacting with conventional allosteric sites on the extracellular surface of the receptor and orthosteric pocket, antibodies are primed to have highly specific allosteric effects^47–50^. The relatively shallow binding modes of AT118-H and AT118-L in AT1R’s orthosteric pocket allows for some, but not all, small-molecule antagonists to simultaneously bind. Small changes to the chemical structure of the antagonists and single amino acid substitutions in AT118 variants drastically alter ligand selectivity (Fig. 5b, 5c, 5e, 5f, Supplementary Data Fig. 5c). In the case of AT118, the probe-dependent allosteric behavior was readily encoded through our ligand-directed nanobody selection strategy, where selected nanobodies displaced the low affinity peptide ligand TRV055 but did not bind to olmesartan bound AT1R^13^. Similar engineering strategies could be readily applied to obtain highly selective probe-dependent allosteric nanobodies for additional receptor families. These molecules could be highly desirable for receptors that bind multiple endogenous ligands, a common feature in GPCR biology. For example, for chemokine and melanocortin receptors, multiple ligands control distinct physiological responses^51,52^. Probe-dependent nanobodies could enhance the action of one ligand over another to achieve the desired receptor biology. Similarly, probe-dependent nanobodies could potentiate the effects of synthetic biased chemical ligands that are designed to engage a subset of signaling outputs^53^. This cooperativity between small molecule and antibody binding adds an increased layer of selectivity to the rich pharmacological potential of nanobodies.

We found that receptor binding and probe-dependence are influenced by separate regions in AT118 family members. Similar binding modes are seen by an allosteric orexin 2 receptor (OX2R) nanobody and apelin receptor (APJ) nanobody antagonist^49,54^. Intriguingly, the APJ nanobody antagonist can be engineered from an antagonist to an agonist through a single amino acid insertion in CDR3^54^. The binding mode of these AT1R, APJ, and OX2R nanobodies is reminiscent of the “message-address” binding site model for opioid receptor ligands, where the address portion of the ligand directs receptor binding and the message region controls pharmacological output (Fig. 6)^55^. This bitopic ligand binding mode combined with the genetic encodability of antibodies presents a readily evolvable platform to obtain a wide range of highly selective GPCR ligands without the need for chemical synthesis.

**Fig 6.**
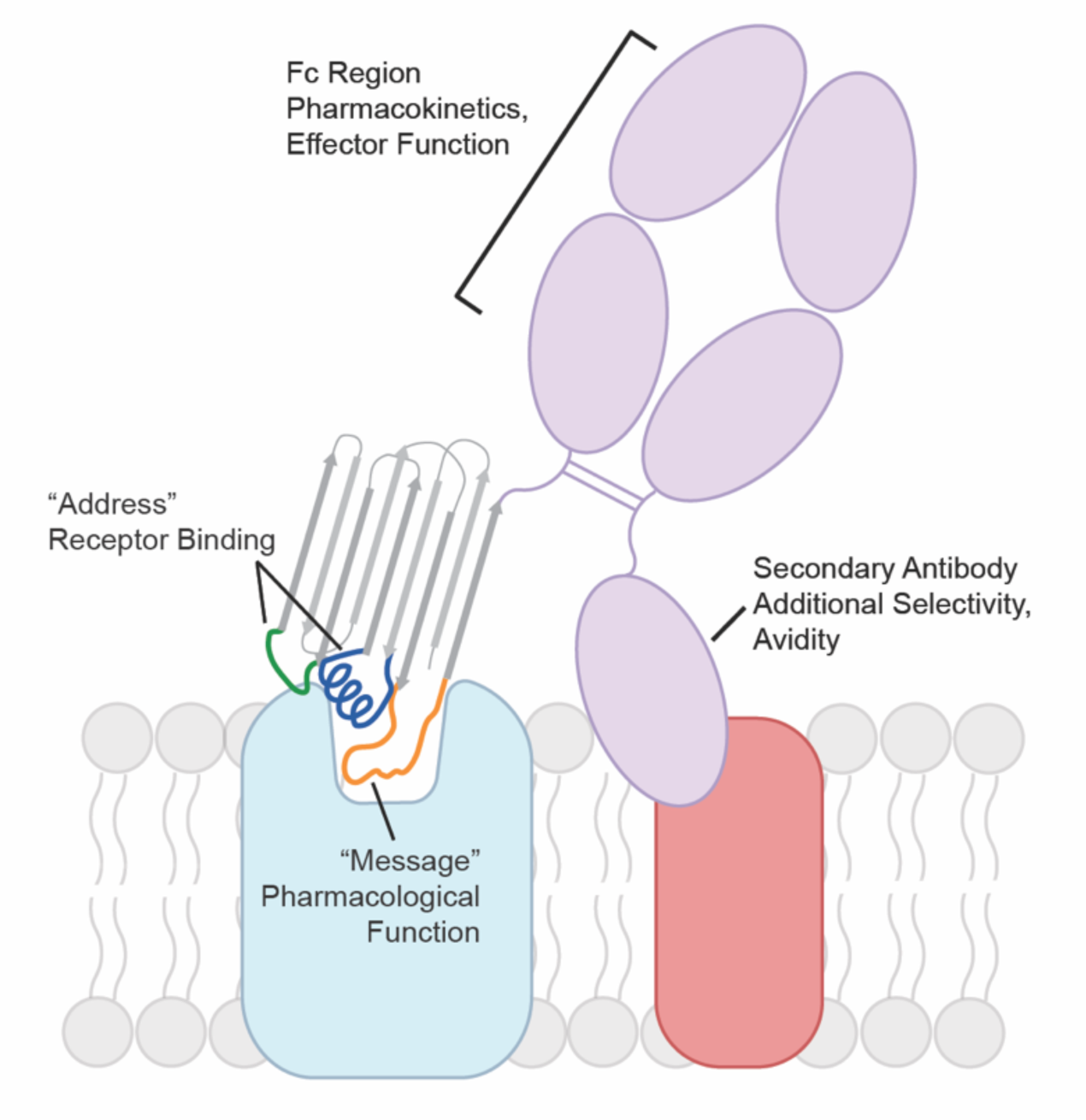
Antibody GPCR binding model. Nanobodies and other antibody fragments can adopt modular binding modes where one region mediates GPCR binding and another influences pharmacological function. The GPCR binding moiety can be formatted into a conventional antibody increasing avidity for the target GPCR or combined with secondary antibodies that recognize a tissue specific marker in a bispecific format. Pharmacokinetics can be further tuned by the antibodies constant Fc region.

In summary, through our characterization of the AT118 family of AT1R nanobody antagonists, we have demonstrated that the modular nature of antibodies facilitates fine-tuning of pharmacokinetic properties to produce ligands that target even well-established therapeutic targets in distinct ways. Our engineered AT118-L Fc fusion proteins elude placental transport and could provide an avenue to selectively target maternal AT1R and determine if the renin-angiotensin system can be modulated during pregnancy to safely regulate blood pressure. Antibody fragments are an untapped pool of pharmacological tools to directly modulate GPCR function. Combining such molecules with conventional antibody engineering approaches, such as bispecifics and altering immune effector functions (Fig. 6), will expand our ability to modulate GPCR function in ways that are not readily accessible to small molecules to ultimately provide new therapeutics and tools for interrogating GPCR function in complex physiological systems.

## Materials and Methods

### Assessing non-specific binding on yeast

Nanobody expression was induced by culturing in -trp + galactose (gal) media for 36-48 hours. 1.5E5 yeast cells were washed with polyspecificity reagent (PSR) staining buffer (20 mM Hepes pH 7.5, 100 mM NaCl, 0.1% n-dodecyl-β-D-maltoside (DDM), 0.01% CHS, 0.05% BSA, 5 mM CaCl2, 10 mM maltose). Cells were stained with 10% *Sf9* insect cell membrane polyspecificity reagent in PSR staining buffer (100 μL volume) at 4 °C for 25 minutes^16,56^. Cells were washed with PSR staining buffer and subsequently stained with 1:200 (v/v) of 1 mg/mL ⍺-HA-AlexaFluor 647 and 1:100 of 0.5 mg/mL Streptavidin-AlexaFluor 488 (100 μL volume) for 15 minutes at 4 °C. Cells were washed and resuspended in PSR staining buffer. Cells, gated for nanobody expression, were analyzed for non-specific binding on a CytoFLEX flow cytometer.

### Targeted library construction

The library of AT118-H variants was constructed by two-step-overlap-extension PCR. A set of ten primers (P94-P103) encoding mixed codons at seventeen amino acid positions was mixed in equimolar ratios (Integrated DNA Technologies, Supplementary Table 6). The mixed primer stock (25-50 nM final concentration) was amplified with Phusion polymerase to create the full length nanobody library DNA. AT118-H library DNA was successively amplified and homology arms corresponding to the pYDS649 plasmid encoding nourseothricin (nat) resistance (pYDS2.0) were added with primers P104-pYDSRev1, pYDSFwd2-pYDSRev2, pYDSFwd3-pYDSRev2. 37.5 μg of YDS2.0 digested with NheI-HF and BamHI-HF (New England Biolabs) and 187.5 μg nanobody insert DNA were transformed into 7.5E9 BJ5465 yeast with a 96-well ECM 830 Electroporator (BTX-Harvard Apparatus). Yeast were recovered in YPAD for 1.5 hours. Transformed cells containing the nanobody plasmid were selected with the addition of nat. Dilutions of transformed yeast were plate on YPAD + nat as single colonies to estimate the library diversity.

### Nanobody selection

5E9 AT118-H yeast library cells were washed in selection buffer 1 (20 mM Hepes pH 7.5, 150 mM NaCl, 2.8 mM CaCl_2_, 0.05% LMNG, 0.005% CHS, 0.1% BSA, 0.2% maltose) and incubated with 1 μM M1 ⍺-FLAG-FITC and a 1:10 (v/v) dilution of anti-FITC magnetic microbeads (Miltenyi) in a 5 mL total staining volume for 40 min at 4 °C. Cells were pelleted and resuspended in 5 mL of selection buffer and applied to a LD column (Miltenyi) to remove yeast expressing nanobodies that bind to M1 ⍺-FLAG-FITC or magnetic beads. After preclearing, the yeast was incubated with 100 nM FLAG-AT1R and 75 nM M1 ⍺-FLAG-FITC for 1 hr at 4° C in selection buffer (5 mL total volume). Cells were pelleted, and resuspended in 5 mL selection buffer with 10% (v/v) anti-FITC magnetic beads (Miltenyi) and incubated for 20 min at 4 °C. Cells were pelleted, washed, and resuspended in selection buffer and applied to an LS column (Miltenyi). 7E6 yeast were eluted from the column.

Yeast from the initial selection round were cultured and 1E8 yeast cells were subjected to a second round of MACS following the same protocol as above with the following modifications. The yeast pool was depleted for binders to both M1 ⍺-FLAG-FITC (200 nM) and M1 ⍺-FLAG-AlexaFluor647 (1 μM) with anti-FITC and anti-AlexaFluor647 magnetic beads. After preclearing, the cells were stained with 50 nM AT1R and 37.5 nM M1 ⍺-FLAG-FITC with 10% (v/v) fetal bovine serum (FBS) added to the staining solution as a blocking reagent for non-specific binders. 1.6E6 cells were recovered and amplified for additional selection rounds using FACS.

2E7 cells were washed with selection buffer and stained with 10% (v/v) *Sf9* insect cell membrane polyspecificity reagent and 1 μM streptavidin-AlexaFluor647 for 30 min at 4 °C in selection buffer. Yeast was washed with 20 mM Hepes pH 7.5, 150 mM NaCl, 2.8 mM CaCl2, 0.1% LMNG, 0.01% CHS, 0.1% BSA, 0.2% maltose followed by 20 mM Hepes pH 7.5, 150 mM NaCl, 2.8 mM CaCl_2_, 0.01% LMNG, 0.001% CHS, 0.1% BSA, 0.2% maltose to remove DDM detergent introduced with the polyspecificity reagent. Cells were then stained with 100 nM FLAG-AT1R and 75 nM M1 ⍺-FLAG-AlexaFluor488 in selection buffer 2 for 30 min at 4 °C. Cells were washed and 2E4 cells preferentially stained with AT1R and ⍺-FLAG-AlexaFluor488 were sorted on a Sony SH800 cell sorter. Sorted cells were amplified and subjected to a final FACS selection round to enrich for clones displaying high affinity towards AT1R. 2E7 cells were washed with selection buffer and stained with 25 nM FLAG-AT1R conjugated to 22.5 nM M1 ⍺-FLAG-AlexaFluor488 and 1:400 ⍺-HA-AlexaFluor647 to detect nanobody expression for 1 hr at 4 °C. Cells were washed and 1E5 cells with high levels of AT1R binding were sorted on a Sony SH800 cell sorter. After each selection round populations were analyzed for nanobody expression, FLAG-AT1R binding, and non-specific binding through small scale analytical staining reactions.

### Sequence analysis

To analyze sequence variation throughout the selection nanobody sequences were directly amplified from the yeast population through PCR with P209 and P210 primers (Supplementary Table 6) and subjected to Illumina sequencing (Azenta, Amplicon-EZ)^16^. Paired-end reads were joined using NGmerge and subsequently filtered using custom scripts to remove reads lacking the amplification primer sequences or containing insertions/deletions, premature stop codons, or ambiguous codons^57^. Amino acid frequencies at every position in the translated nanobodies were calculated using custom scripts.

### AT1R staining on yeast

Plasmids containing AT118-H variants were transfected into BJ5465 yeast following the PEG/lithium acetate method^58^. Nanobody expression was induced by culturing in -trp + gal media for 36-48 hours. 1.5E5 yeast cells were washed with selection buffer and stained with 100 nM FLAG-AT1R and 35 nM M1 ⍺-FLAG-AlexaFluor 488 and 1:200 (v/v) of 1 mg/mL ⍺-FLAG-AlexaFluor 647 (50 μL volume) at 4 °C for 25 minutes. Cells were washed and resuspended in selection buffer. Cells, gated for nanobody expression, were analyzed for AT1R binding on a CytoFLEX flow cytometer.

### Nanobody expression and purification

AT118 variants were cloned into a pET26b vector with an N-terminal PelB signal sequence for periplasmic expression and a C-terminal V5 epitope followed by a hexahistidine tag. Plasmids were transformed into BL21(DE3) *E. coli* and grown in TB supplemented with 4% (v/v) glycerol and 50 μg/mL kanamycin. Cultures were grown to an OD_600_ of 2 at 37 °C and cooled to 20 °C for 1 hour. Protein expression was induced with 0.2 mM IPTG.

Cell pellets were resuspended in room temperature SET buffer (200 mM Tris pH 8, 500 mM sucrose, 500 μM EDTA) and stirred for 20 min. Resuspended cells were diluted with 2 volumes of cold H_2_O, supplemented with 5 mM MgCl_2_ and benzonase nuclease, and stirred for an hour. Cell debris was pelted by centrifugation (14,000xg, 30 min). 100 mM NaCl was added to the supernatant and stirred for 15 min. The supernatant was filtered and passed over Protein A resin (GoldBio), washed with 10 mM sodium phosphate pH 7.5, 100 mM sodium chloride, and eluted with 100 mM sodium phosphate pH 2.5, 100 mM sodium chloride directly into 2M Hepes pH 8 to rapidly neutralize the eluent. Proteins were dialyzed into HBS + 10% glycerol and flash frozen in liquid nitrogen. The AT118-L MBP fusion protein was purified following the same protocol with the wash, elution, and dialysis buffers supplemented with 2 mM maltose^24^.

Nanobodies used for AT1R binding assays were further purified by nickel affinity chromatography to ensure the C-terminal epitope was intact. The protein A column eluent was passed over Ni-NTA resin, washed with 20 mM Hepes pH 7.5, 150 mM sodium chloride, and eluted with 20 mM Hepes pH 7.4, 150 mM sodium chloride, 200 mM imidazole. Nanobodies were dialyzed into HBS + 10% glycerol and flash frozen in liquid nitrogen. Protein purity was analyzed by SDS-PAGE.

### Nanobody-Fc construct design, expression, and purification

AT118-H followed by a short GSSG linker was cloned into the pFUSE-CHIg-hG1 (InvivoGen) to fuse it to human IgG1 Fc. Antibody effector function was silenced by introducing mutations into the IgG1 Fc to block Fc gamma receptor binding (L234A, L235A) [IgG1 Fc residue numbering is according to the Eu numbering scheme^59^] and complement fixation (P329G). To inhibit FcRn binding, H435A, I253A, and H310A substitutions blocking both antibody recycling and transport of antibodies across the placenta. To generate a non-AT1R binding control nanobody-Fc fusion we introduced F47T^CDR2^ and Y98F^CDR3^ substitutions. Plasmids (750 μg / L of culture) were transfected into Expi293 cells grown in Expi293 media with Fectopro (800 μL / L of culture). 6 mM valproic acid and 0.8% glucose were added after 18-24 hours. Cells were pelleted 5-7 days post transfection and the supernatant was applied to protein A resin. Resin was washed with 10 column volumes of 20 mM Hepes pH 7.4, 150 mM NaCl and eluted with 100 mM glycine pH 2.5 directly into 1 M Hepes pH 8. Purified nanobody-Fc fusion proteins were dialyzed overnight into PBS (10 mM Na_2_HPO_4_, 1.8 mM KH_2_PO_4_, 2.7 mM KCl, 137 mM NaCl). Protein for mouse experiments was diluted to 1 mg/mL and sterile filtered.

### Radioligand binding competition assays

Cell membranes for radioligand binding experiments were prepared from a tetracycline inducible stable human FLAG-AT1R Expi293F cell line^16^. AT1R expression was induced at 2×10^6^ cells/mL with 0.4 μg/mL doxycycline hyclate for 30 hours. Cells were pelleted and washed with cold HBS (20 mM Hepes pH 7.4, 150 mM NaCl). Cells were resuspended in 2.5 mL of 20 mM Tris pH 7.4 per gram of cell pellet with a protease inhibitor tablet and lysed by Dounce homogenization (100x). Membranes were pelleted by centrifugation at 50,000 x g for 20 min and resuspended in 2.5 mL of 50 mM Tris pH 7.4, 12.5 mM MgCl_2_, 150 mM NaCl, 0.2% BSA + protease inhibitor table by Dounce homogenization, flash frozen in liquid nitrogen, and stored at −80°C.

Membranes were incubated with nanobody variants and 2 nM [^3^H]-olmesartan (American Radiolabeled Chemicals) in 50 mM Tris pH 7.4, 12.5 mM MgCl2, 150 mM NaCl, 0.2% BSA in a 200 μL reaction volume for 90 minutes at room temperature. Reactions were harvested on a GF/B filter soaked in water on a 96-well Brandel harvester and washed three times with cold water.

### G⍺_q_ signaling assay

Expi293F cells stably expressing a tetracycline inducible wild-type human FLAG-AT1R were diluted to 2 x 10^6^ cells/mL and induced with 0.4 μg/mL doxycycline hyclate for 18-20 hours. 2 x 10^4^ cells were plated into a low-volume 96-well plate, treated with 5 μM of nanobody or nanobody-Fc variant for 30 min at 37 °C, and stimulated with AngII for 1 hr at 37 °C. IP1 was detected with the IP-One Gq kit (CisBio) and read on a SpectraMax M5e plate reader (Molecular Devices).

### Placental transfer assay

The Boston Children’s Hospital IACUC approved all experiments involving mice for this study. Timed pregnant CD-1 mice (Charles River Laboratories) at embryonic day 15.5 (E15.5) were treated with one dose of theAT118-L-Fc fusion proteins (5 mg/kg) via intraperitoneal (IP) injection. Animals were housed in a temperature-and humidity-controlled room with 12-hr light / 12-hr dark cycle and had free access to food and water. Maternal and fetal blood were collected 48 hours later from the heart. Serum samples were prepared by letting whole blood samples coagulate at room temperature for 30 minutes followed by centrifugation at 5000xg at room temperature for 10 minutes to collect the supernatant (serum). Maternal and fetal serum was analyzed for Fc-fusion proteins using a Human-Fc ELISA assay (SydLabs). Serum concentrations were determined by generating a standard curve for each construct.

### AT1R construct design

All AT1R constructs were cloned into a pCDNA3.1-Zeo-tetO vector following an N-terminal hemagglutinin signal sequence (Supplementary Table 3)^60^. Wild-type AT1R was fused to a N-terminal FLAG tag for affinity purification^25^. To boost receptor expression for structural studies, AT118-H was inserted after the hemagglutinin signal sequence followed by a short linker (LEGGSG) and protein-C epitope sequence and directly fused to the N-terminus of AT1R. The receptor was truncated at residue 320 to remove the unstructured C-terminus and followed by a 3C protease site and Rho 1D4 epitope tag. Two fusion constructs were cloned to enhance the size of the complex. To generate an AT1R fusion capable of binding alpacaized Nb6, a nanobody that binds ICL3 of the kappa opioid receptor, AT1R residues 218-243 were replaced with 249-277 of the human kappa opioid receptor^23^. To generate a rigid AT1R-BRIL fusion AT1R residues 227-233 were replaced with thermostabilized apocytochrome b_562_RIL (BRIL) followed by seven amino acids of helix 6 of the human A_2A_ adenosine receptor and three amino acids from helix 6 of the human frizzled 5 receptor^61^.

### Quantifying receptor expression

1.8 mL Expi293 Tet-R cells grown to 3E6 cells/mL in Expi293 media were plated in a 6 well plate. 1.5 μg of receptor plasmid (pCDNA3.1 tet/zeo) was transfected with FectoPro (1.6 μL). Cells were enhanced with 6 mM valproic acid and 0.8% glucose 16-20 hours after transfection. Receptor expression was induced with 4 μg/mL doxycycline hyclate and 5 mM sodium butyrate. After 24 hours cells were pelleted (500 x g, 4 min), washed with PBS, and fixed with 0.01% (v/v) formaldehyde in PBS. Cells were washed twice with PBS and permeabilized with 0.5% Tween20 (v/v) in PBS. Cells were washed twice with PBS followed by HBS + 0.1% BSA (w/v) Cells were stained with 1:1000 (v/v) of ⍺-1D4-AlexaFluor488 antibody in HBS + 0.1% BSA. Washed with HBS + 0.1% BSA and analyzed on a CytoFLEX flow cytometer.

### AT1R expression and purification

AT1R plasmids (750 μg / L of culture) were transfected into Expi293 cells with a stably integrated tetracycline repressor grown in Expi293 media with Fectopro (800 μL / L of culture). 6 mM valproic acid and 0.8% glucose were added after 18-24 hours. 5 μM kifunensine was added to cultures transfected with constructs for cryo-EM. Two days after transfection cells were induced with 0.4 μg/mL doxycycline and 5 mM sodium butyrate. For constructs lacking an N-terminal nanobody fusion, 5 μM losartan was included during induction. Pellets were harvested 30 hours post induction.

Frozen cell pellets were resuspended in room-temperature hypotonic lysis buffer (10 mL 10 mM Tris pH 7.4, 2 mM EDTA, 10 mM MgCl_2_ / g cell pellet), with benzonase nuclease (Sigma Aldrich), Pierce protease inhibitor tablets (ThermoFisher). Cells were pelleted (50,000 x g, 15 min) and resuspended in (10 mL / g initial cell pellet) cold solubilization buffer (20 mM Hepes pH 7.4, 500 mM NaCl, 0.5% lauryl maltose neopentyl glycol (LMNG), 0.05% cholesterol hemisuccinate (CHS), 10 mM MgCl_2_, benzonase nuclease, Pierce protease inhibitor tablets) and disrupted with a Dounce homogenizer. Resuspended cells were stirred for two hours at 4 °C. Insoluble material was pelleted by centrifugation (50,000 x g, 30 min). The supernatant was filtered through a glass fiber filter, supplemented with 2 mM CaCl_2_ and passed over M1-⍺FLAG or HPC-4-⍺Protein C resin. The resin was washed with 20 column volumes of wash buffer (20 mM Hepes pH 7.4, 500 mM NaCl, 0.01% LMNG, 0.001% CHS, 2 mM CaCl_2_) and eluted with 20 mM Hepes pH 7.4, 500 mM NaCl, 0.01% LMNG, 0.001% CHS, 5 mM EDTA with 0.2 mg/mL FLAG (DYKDDDK) or Protein C peptide (EDQVDPRLIDGK). The receptor was further purified and buffer exchanged by size exclusion chromatography on a Superdex S200 (10/300) Increase column in 20 mM Hepes pH 7.4, 100 mM NaCl, 0.01% LMNG, 0.001% CHS. For receptors lacking the N-terminal nanobody fusion, 5 μM losartan was included during lysis and solubilization.

### Fab expression and purification

BAG2 (anti-BRIL Fab) and NabFab (anti-nanobody Fab) heavy chains were cloned into pTarget and light chains were cloned into pD2610^21,22^. Light and heavy chain plasmids (375 μg / L of culture) were transfected into Expi293 cells grown in Expi293 media with Fectopro (800 μL / L of culture). 6 mM valproic acid and 0.8% glucose were added after 18-24 hours. Cells were pelleted 5-7 days post transfection and the supernatant was applied to CaptureSelect IgG-CH1 resin. Resin was washed with 10 column volumes of 20 mM Hepes pH 7.4, 150 mM NaCl and eluted with 100 mM glycine pH 2.5 directly into 1 M Hepes pH 8 (5:10 v/v ratio). Fab was dialyzed into 20 mM Hepes pH 7.4, 150 mM NaCl, 10% glycerol and flash frozen until use.

### Cryo-EM sample preparation

AT118-H-proteinC-AT1R-BRIL was mixed with 1.4-fold molar excess of Anti-BRIL Fab, and 2-fold molar excess of anti-Fab nanobody and incubated on ice for one hour. The protein mixture was supplemented with 2 mM CaCl_2_ loaded onto 1 mL of HPC4-⍺Protein C resin and washed with 6 CV of 20 mM Hepes pH 7.4, 100 mM NaCl, 0.01% LMNG, 0.001% CHS, 2 mM CaCl_2_. The column was then washed with 20 CV of 20 mM Hepes pH 7.4, 100 mM NaCl, 0.05% glyco-diosgenin (GDN), 0.005% CHS, 2 mM CaCl_2_ to exchange detergent. The complex was eluted with 20 mM Hepes pH 7.4, 100 mM NaCl, 0.05% GDN, 0.005% CHS + 5 mM EDTA + 0.2 mg/mL Protein C peptide. The complex was concentrated in a 100 kDa cutoff Amicon-Ultra centrifugal filter, supplemented with 2-fold molar excess of anti-Fab nanobody, and further purified by size exclusion chromatography (Superdex S200 Increase 10/300) in 20 mM Hepes pH 7.4, 100 mM NaCl, 0.05% GDN, 0.005% CHS. Peak fractions were concentrated to 9.2 mg/mL

FLAG-AT1R-BRIL was mixed with a 3-fold molar excess of AT118-L, a 1.4-fold molar excess of Anti-BRIL Fab, and 2-fold molar excess of anti-Fab nanobody and incubated on ice for one hour. The protein mixture was supplemented with 2 mM CaCl_2_ loaded onto 1 mL of M1-⍺FLAG resin and washed with 6 CV of 20 mM Hepes pH 7.4, 100 mM NaCl, 0.01% LMNG, 0.001% CHS, 2 mM CaCl_2_. The column was then washed with 20 CV of 20 mM Hepes pH 7.4, 100 mM NaCl, 0.05% GDN, 0.005% CHS, 2 mM CaCl_2_ to exchange detergent. The complex was eluted with 20 mM Hepes pH 7.4, 100 mM NaCl, 0.05% GDN, 0.005% CHS + 5 mM EDTA + 0.2 mg/mL FLAG peptide. The complex was concentrated in a 100 kDa cutoff Amicon-Ultra centrifugal filter, supplemented with 2-fold molar excess of anti-Fab nanobody, and further purified by size exclusion chromatography (Superdex S200 Increase 10/300) in 20 mM Hepes pH 7.4, 100 mM NaCl, 0.05% GDN, 0.005% CHS. Peak fractions were concentrated to 7 mg/mL.

3 μL sample was applied to glow-discharged UltrAuFoil 300 mesh grids, 1.2 μm diameter /1.3 μm spacing (TedPella) at 20 °C with 100% humidity. Grids were blotted for 4-6 seconds with blotforce of 15 on a Vitrobot Mark IV Vitrobot (ThermoFisher) and directly plunged into liquid-nitrogen-cooled liquid ethane.

### Cryo-EM data collection, processing, and model building

Cryo-EM data were collected on a 300 kV Titan Krios G3i microscope (Thermo Fisher) with a K3 direct electron detector (Gatan) and a GIF quantum energy filter (20 eV) (Gatan) in counted mode at the Harvard Cryo-Electron Microscopy Center for Structural Biology. A single movie was acquired in the center of nine holes per stage position with the SerialEM data collection software^62^. Data collected parameters are listed in Supplementary Table 4.

All data processing steps were performed in CryoSPARC (Supplementary Data Fig. 4 and 5)^63^. Movies underwent patch motion correction, dose-weighting, and patch contrast transfer function (CTF) estimation. Particles were picked with the blob-picker function. A subset of particles resembling our samples were identified through 2D classification from a preliminary data collection and used to generate *ab initio* reconstructions. All particles were then classified through successive rounds of heterogenous refinement followed by non-uniform refinement with global CTF refinement. A local mask excluding the CH1 and CL domains of the anti-BRIL Fab and anti-Fab nanobody was applied for local refinement. The model was built in Coot and refined with Phenix real space refinement^64,65^. In regions with strong backbone density, but weak density for side chains, side chains were modeled in their most common rotamer position that did not clash with the model. Notably, no density was observed for the first β-strand of AT118-H in the AT118-H-AT1R fusion protein. We sequenced the protein by Edman degradation and confirmed that this region is proteolyzed during expression and purification but does not distort the overall nanobody fold. Figures were prepared in UCSF Chimera and Pymol^66,67^. All structural biology software was compiled by SBGrid^68^.

### AT118 AT1R binding assays

To test the ability of AT1R variants to bind AT118-H with a C-terminal V5 tag 900 uL Expi293 TetR cells were plated in a 12 well plate at 3E6 in Expi293 media. 750 μg of pCDNA3.1 tet/zeo plasmid containing FLAG-AT1R variants were transfected into the cells with FectoPro (0.8 μL). Cells were enhanced with 6 mM valproic acid and 0.8% glucose 16-20 hours after transfection. Receptor expression was induced with 4 μg/mL doxycycline hyclate and 5 mM sodium butyrate. After 18 hours 1.5E5 cells were pelleted (300xg, 5 min), washed with flow assay buffer (20 mM Hepes pH 7.4, 150 mM NaCl, + 0.1% BSA), and stained with 20 nM AT118-H (1 mL staining volume). Cells were washed twice with flow assay buffer, stained with 100 nM M1-⍺FLAG-AlexaFluor488 and 1:500 (v/v) ⍺V5-AlexaFluor647 in flow assay buffer + 2 mM CaCl_2_. Cells were washed once and resuspended with flow assay buffer + 2 mM CaCl_2_.

To analyze the probe dependence of AT118-H and AT118-L a stable FLAG-AT1R Expi293F cell line was induced at 2E6 cells/mL with 0.4 μg/mL doxycycline hyclate for 18 hours. 1.5E6 cells were washed with flow assay buffer and incubated with 10 μM of various ligands for 20 min at 4 °C in a 50 μL volume. 50 μL of 40 nM V5 epitope tagged AT118-H or AT118-L was added and the samples were incubated for an hour at 4 °C with shaking at 30 rpm. Cells were washed twice with flow assay buffer and stained with 100 nM M1-⍺FLAG-AlexaFluor488, 1:500 (v/v) anti-V5-AlexaFluor647 antibody (ThermoFisher), 2 mM CaCl_2_, and 5 μM of the appropriate ligand for 20 min at 4 °C. C ells were washed and resuspended in flow assay buffer supplemented with 2 mM CaCl_2_. Purity and mass of olmesartan derivatives were confirmed by HPLC and HR-MS (Supplementary Table 5).

For saturation binding experiments a stable FLAG-AT1R Expi293F cell line was induced at 2E6 cells/mL with 0.4 μg/mL doxycycline hyclate for 18 hours. 2.8E5 cells were plated and stained with 0-1 μM of each AT118-V5 tag or AT118 Fc variant in flow assay buffer (100 µL reaction volumes) for 1 hour at 4°C with shaking at 30 rpm. Cells were washed 2 times with flow assay buffer and subsequently stained with 100 nM of M1-⍺FLAG-AlexaFluor488 and 1:500 (v/v) of Alexafluor 647 conjugated anti-V5 antibody (ThermoFisher) (for AT118-H or AT118-L) or 1:1000 (v/v) Alexafluor 647 conjugated Anti-Human IgG Fc (Biolegend) (for AT118-L Fc Fusion proteins) in flow assay buffer + 2 mM CaCl_2_ (100 µL reaction volumes) for 20 minutes at 4 °C. Cells were washed once and resuspended in flow assay buffer + 2 mM CaCl_2_. Levels of AT118 or AT118-Fc variant binding was assessed on a CytoFLEX Beckman Coulter. Single cells were gated for receptor expression and analyzed with CytoFLEX software and GraphPad Prism 9.5.1.

## Acknowledgements

We thank R. Lefkowitz for allowing preliminary pharmacological experiments to be performed in his lab, M. Bao and X. Zhou for critical reading of the manuscript, A. Kossiakoff for the BAG2 anti-BRIL Fab, and R. Walsh and M. Mayer for assistance and advice during cryo-EM data collection at the Harvard Center for Cryo-Electron Microscopy at Harvard Medical School. The SBGrid consortium provided computation support of structural biology software. This work was funded by a Merck Fellowship from the Helen Hay Whitney Foundation (M.A.S.); NIH Grants DP5OD021345 (A.C.K.), R21HD101596 (A.C.K.), R01CA260415 (A.C.K.), R01NS088566 (M.K.L); the Vallee Foundation (A.C.K.); the Smith Family Foundation (A.C.K.); the Pew Charitable Trusts (L.M.W.); the Whitehead Foundation (L.M.W.); the New York Stem Cell Foundation (M.K.L); and the William Randolph Hearst Fellowship (H.X.).

## Author Contributions

M.A.S., S.M.S., S.R., D.P.S., L.M.W., C.M., and A.C.K., designed experiments. M.A.S., L.M.W., S.M.S., M.S.A.G., G.R.N., H.X., P.S., and J.K. performed experiments. M.A.S., L.M.W., S.M.S., M.S.A.G., S.R., and J.D.H analyzed data. M.K.L. and A.C.K. supervised research. M.A.S. wrote the paper with input from all authors.

## Data and Materials Availability

The cryo-EM model and maps were deposited under the following accession numbers AT118-H AT1R complex PDB: 8TH3, EMDB: 41248; AT118-L AT1R Complex PDB: 8TH4, EMDB: 41249. Requests for materials should be addressed to Andrew C. Kruse.

## Code Availability

Custom code for analyzing sequencing data is available upon request.

