## Supplementary information for "Antibodies Expand the Scope of Angiotensin Receptor Pharmacology"

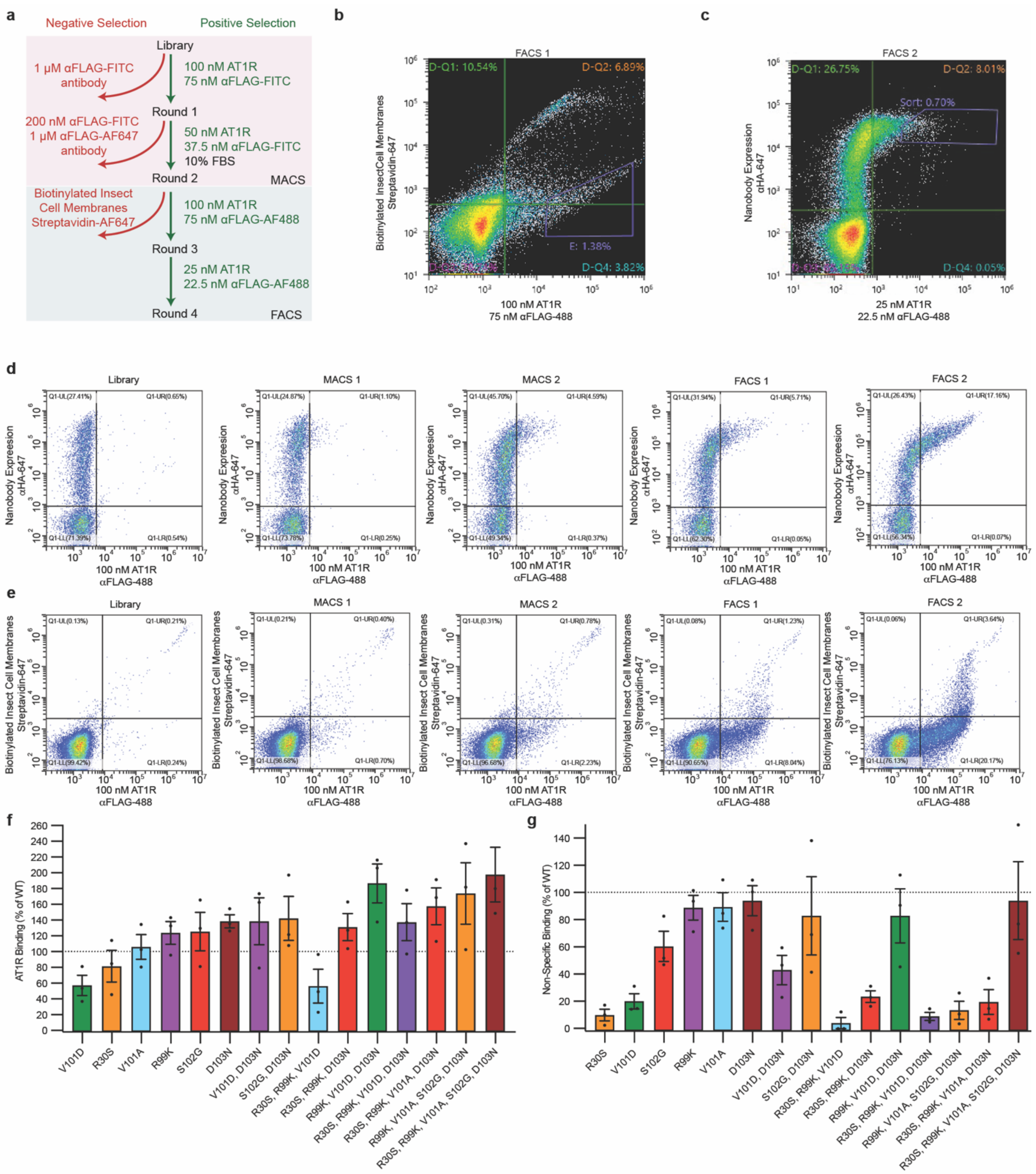

**Supplementary Figure 1. Engineering of high affinity AT1R nanobody antagonist with low non-specific binding.** a) Flowchart of nanobody selection. AT1R binders were enriched through two rounds of magnetic-activated cell sorting (MACS). Fluorescence-activated cell sorting (FACS) was used to isolate clone with low polyreactivity. A final FACS step enriched high-affinity AT1R binders. b) FACS round 1 plot. 1.38% of the population containing high-affinity AT1R binders with reduced polyspecificity were collected. c) FACS round 2 plot 0.7% of the population was collected containing high affinity AT1R binders. d) Binding of yeast-display library to FLAG-AT1R throughout each selection round. e) Distribution of AT1R-binding and polyreactive nanobodies in the yeast-display library throughout the selection process. f) Binding of AT118-H variants displayed on yeast to detergent solubilized AT1R. Error bars represent mean ± standard error from three experiments. g) Non-specific binding AT118-H variants displayed on yeast to biotinylated insect cell membrane polyspecificity reagent. Error bars represent mean ± standard error from three experiments.

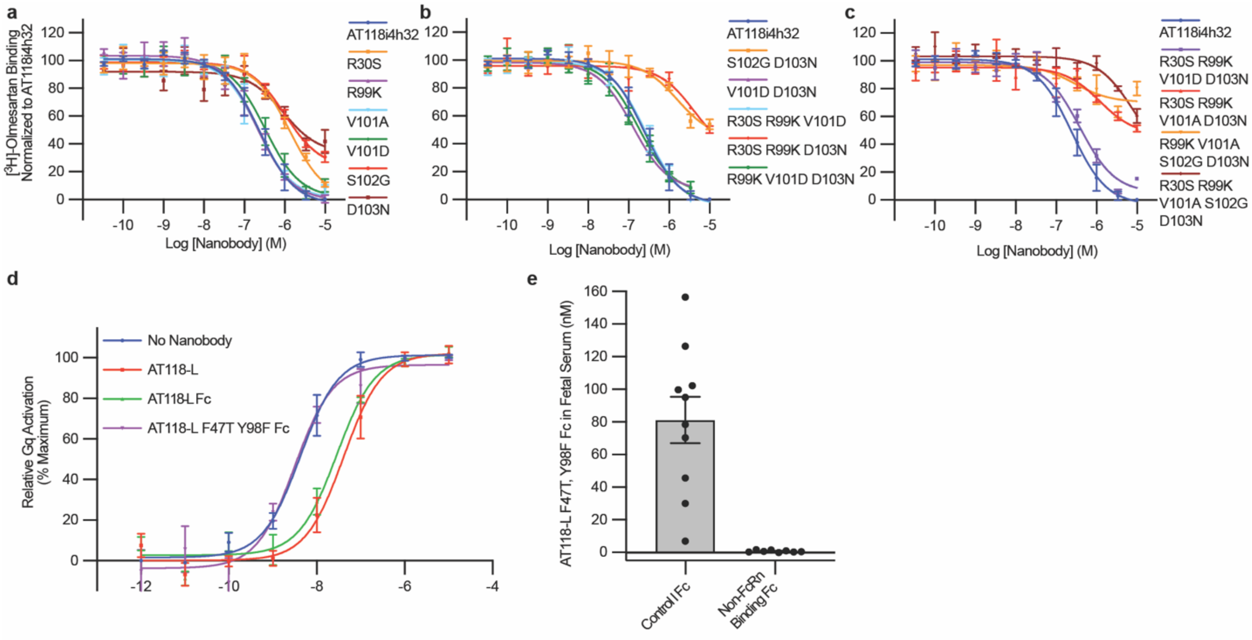

**Supplementary Data Figure 2. Effects of AT118-L and AT118-L-Fc fusion proteins on AT1R signaling.** a-c) $^{3}H$-olmesartan competition experiments in Expi293 cell membranes containing AT1R with purified AT118-H variants. Variants containing D103N and S102G fail to displace olmesartan. The addition of V101D to D103N rescues the loss in pharmacological function. Error bars represent mean ± standard error from three experiments. Error is too small to be displayed if no bar is present. d) AT118-L Fc fusion proteins (pMAS493, Supplementary Table 1) suppress $G\alpha_q$ signaling (IP1 accumulation), whereas AT118-L F47T Y98F (pMAS513, Supplementary Table 1), the non-AT1R binding control, Fc fusion protein has no effect. Error bars represent mean ± standard error from three experiments. e) Accumulation of AT118-L F47T Y98F Fc fusion protein, that does not bind AT1R, in fetal serum. Error bars represent mean ± standard error from nine embryos from three separate litters for the control Fc (pMAS512, Supplementary Table 1) and eight embryos from two litters for the engineered Non-FcRn binding Fc (pMAS513, Supplementary Table 1).

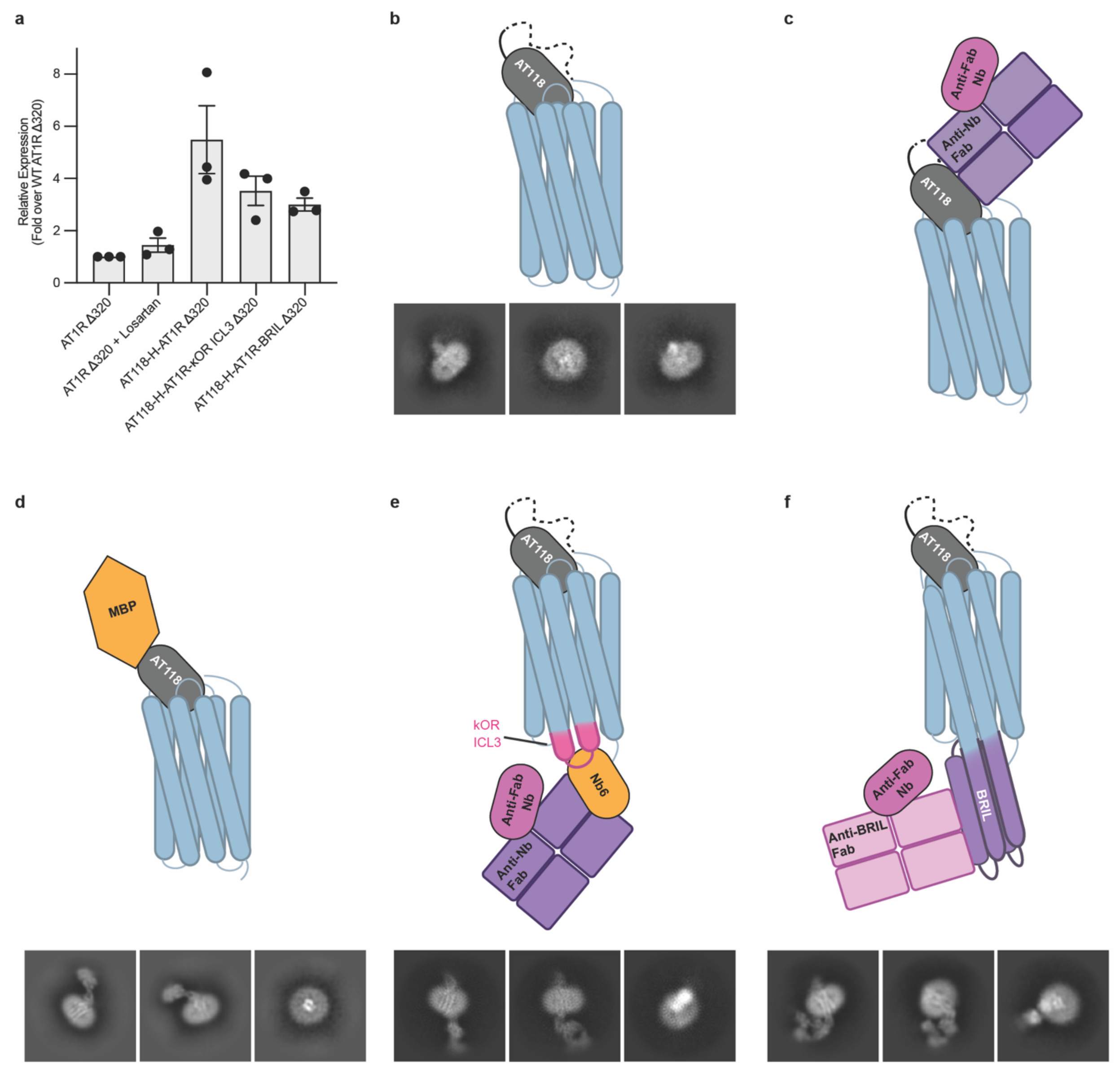

**Supplementary Data Figure 3. Cryo-EM Construct Screening.** a) Fusion of AT118-H to the N-terminus of AT1R enhances total receptor expression. Error bars represent mean ± standard error from three experiments. b-f) Constructs screened for structure determination and representative 2-D class averages. b) AT118-AT1R fusion protein, c) Anti-nanobody Fab fragment bound to free AT118, but not AT118 in complex with AT1R[22]. d) MBP-AT118 in complex with AT1R[24]. e) AT118-AT1R-kappa opioid receptor (κOR) ICL3 fusion protein in complex nanobody 6 with an engineered alpaca framework, which binds κOR ICL3, anti-nanobody Fab, and anti-Fab nanobody[22,23]. f) AT118-AT1R-BRIL fusion protein in complex with anti-BRIL Fab and an anti-Fab nanobody[21].

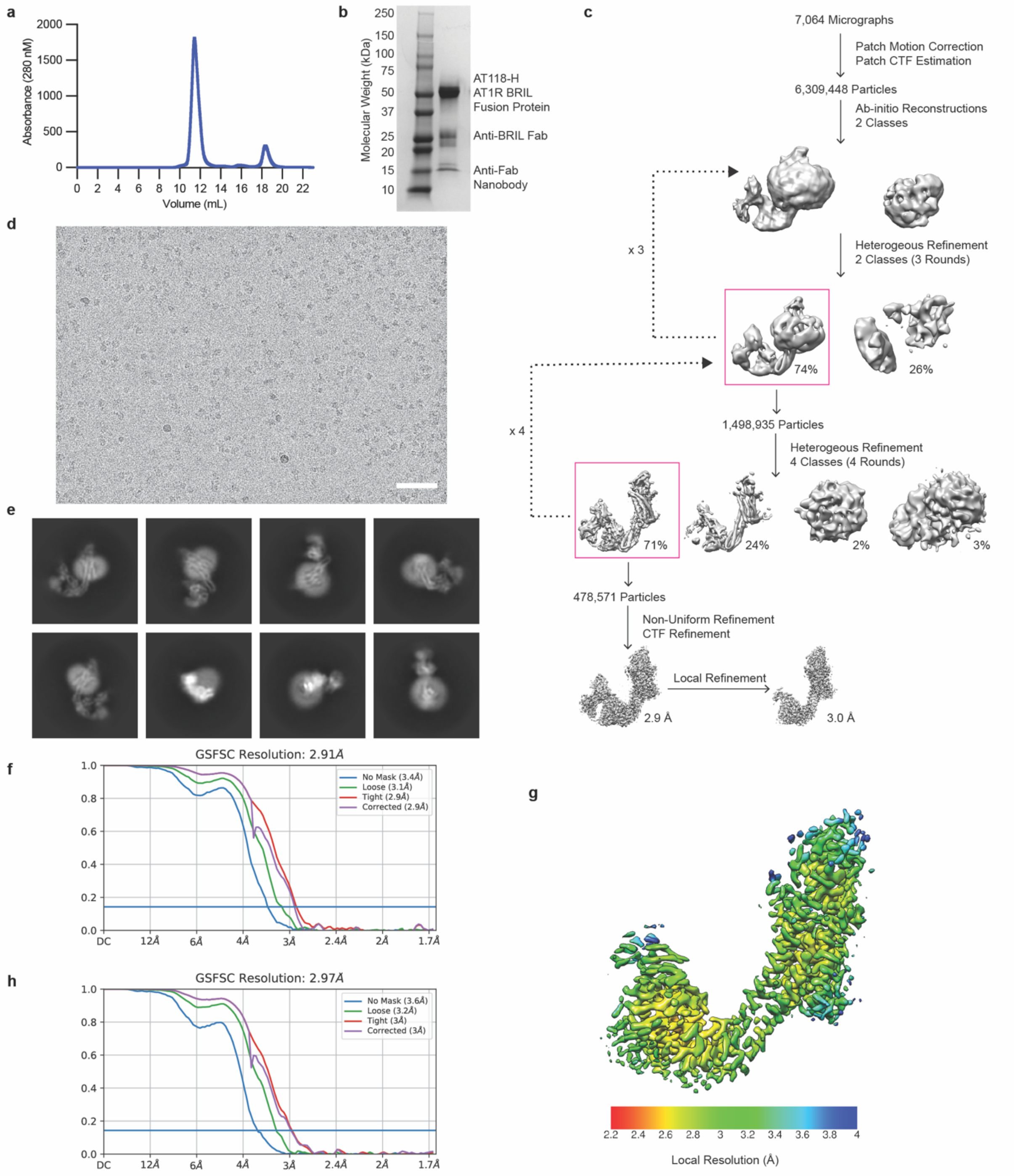

**Supplementary Data Figure 4. AT118-H AT1R Data Processing.** a) Size exclusion trace, b) SDS-PAGE gel under reducing conditions, c) cryo-EM data processing scheme, d) representative micrograph (scale bar = 50 nm), and e) representative 2D class averages of AT118-H-AT1R-BRIL, anti-BRIL Fab, anti-Fab nanobody complex. f) Fourier shell correlation (FSC) used to determine the global map resolution. g) Local resolution estimate after local refinement. h) FSC used to determine locally refined map resolution.

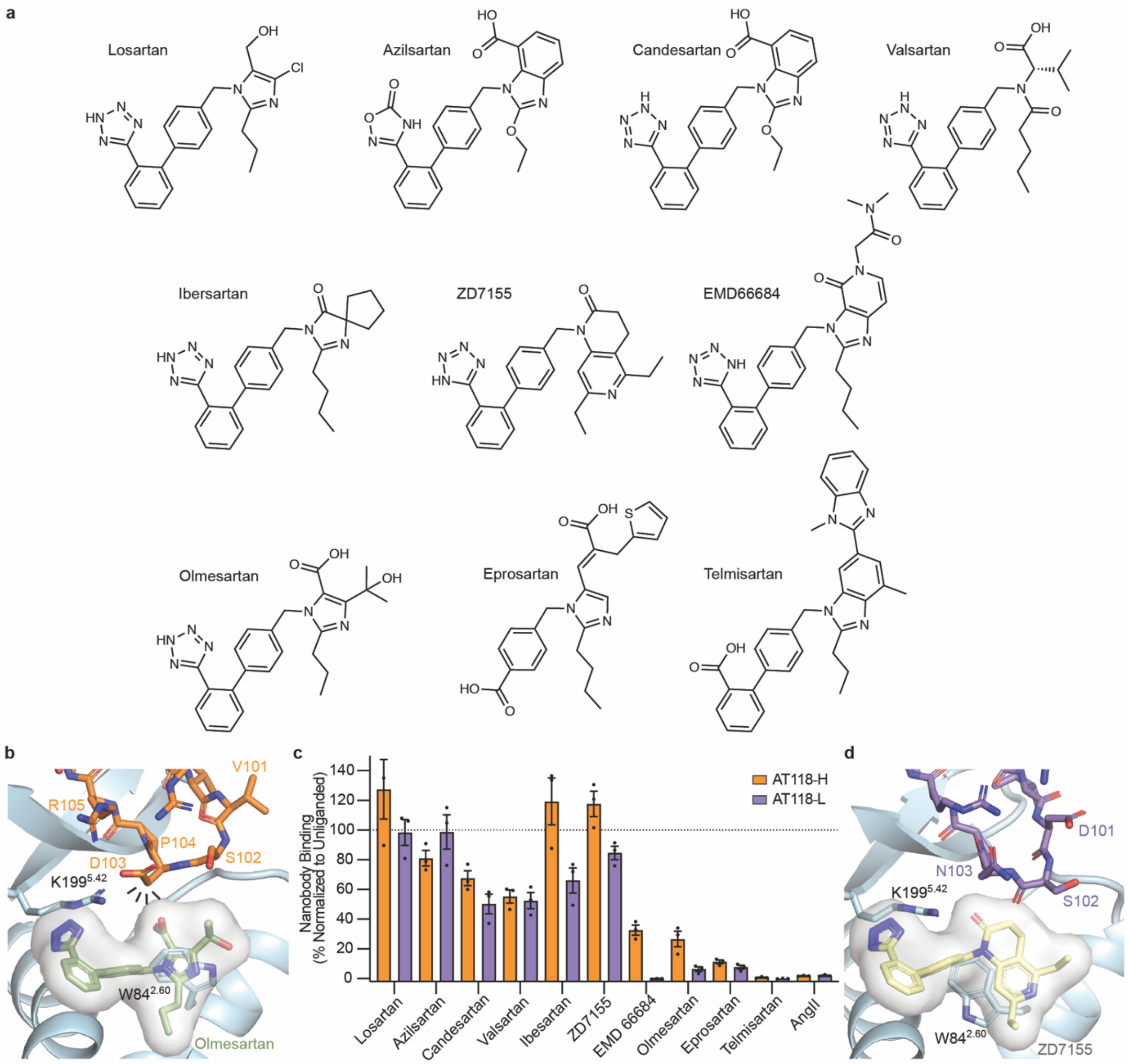

**Supplementary Data Figure 5. Binding of AT118-H and AT118-L with a broad panel of small molecule AT1R antagonists.** a) Molecular structures of small-molecule AT1R antagonists. b) D103$^{CDR3}$ of AT118-H would clash with modeled olmesartan. Weak density for W84$^{2.60}$ is observed in the antagonist binding site in orthosteric pocket of the H-AT1R fusion protein structure. c) Binding of AT118-H (orange) and AT118-L (purple) with a series of small-molecule AT1R antagonists. Error bars represent mean ± standard error from three experiments. d) Binding of AT118-L with modeled ZD7155 (pink sticks, PDB 4YAY[31]).

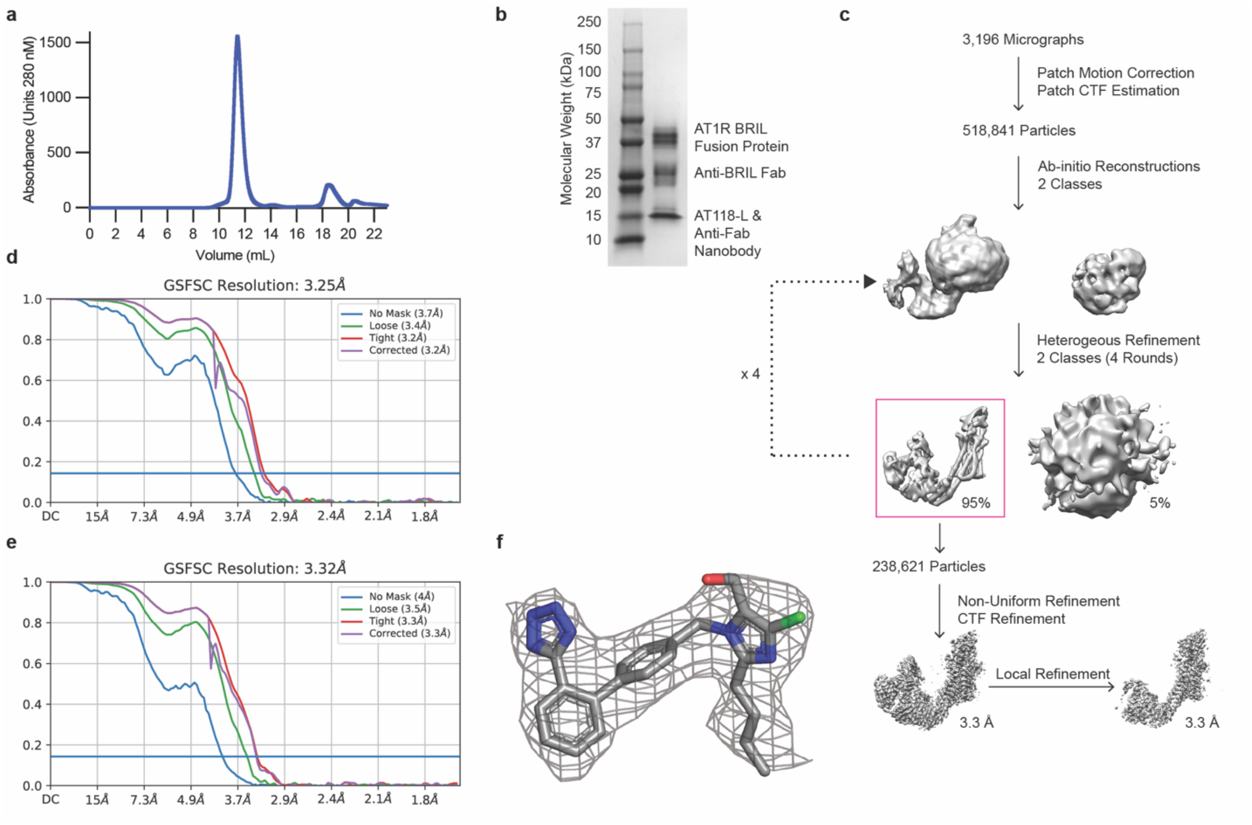

**Supplementary Data Figure 6. AT118-L AT1R Data Processing.** a) Size exclusion trace, b) SDS-PAGE gel under reducing conditions, and c) cryo-EM data processing scheme of AT118-L AT1R-BRIL, anti-BRIL Fab, anti-Fab nanobody complex. d) Fourier shell correlation (FSC) used to determine the global map resolution. e) FSC used to determine locally refined map resolution. f) Experimental density of losartan within the orthosteric binding pocket from locally refined map.

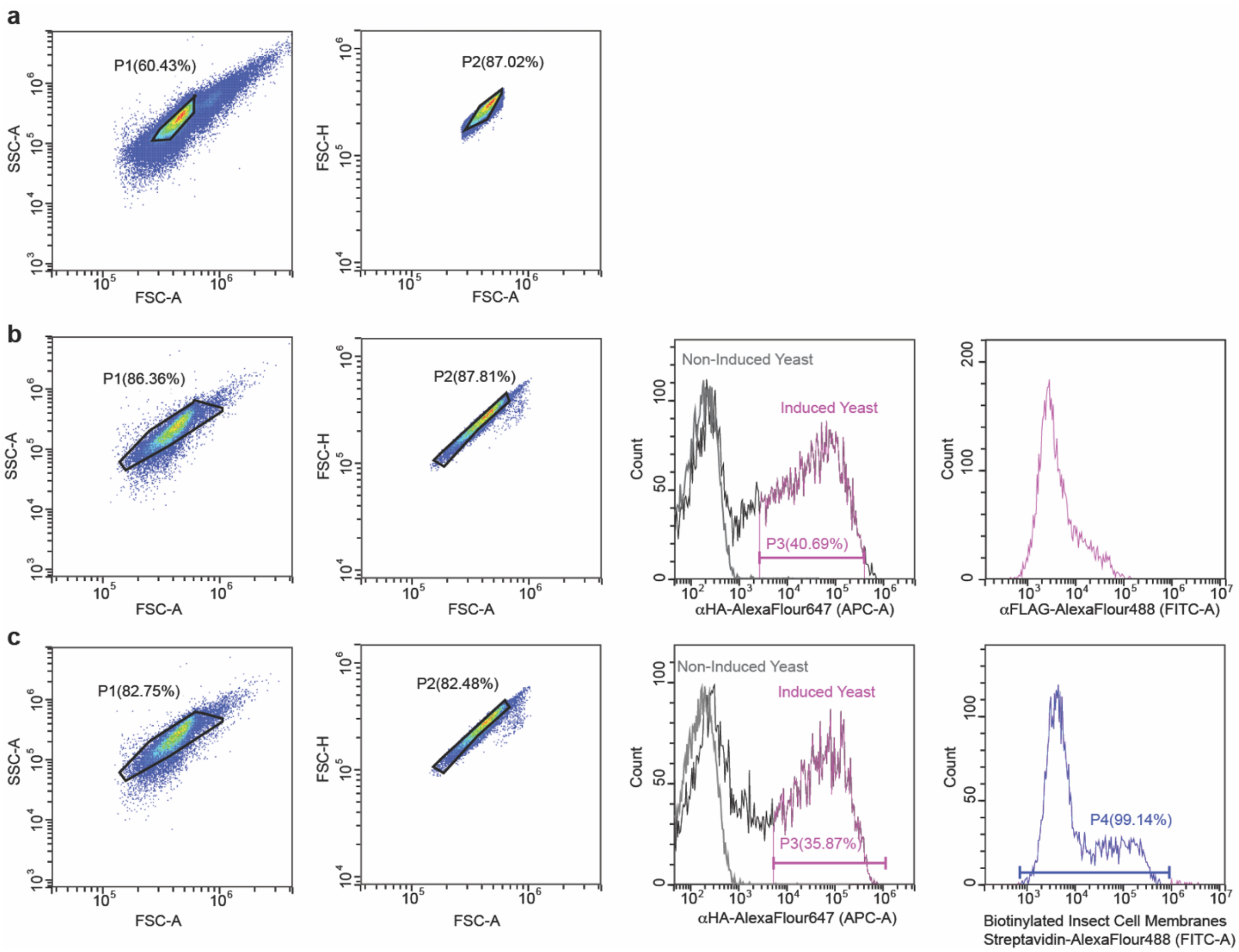

**Supplementary Data Figure 7.** Representative flow cytometry gating of yeast cells. Flow cytometry plots illustrating gating strategies for a) yeast cells displaying nanobodies in Supplementary Data Fig. 1b-e; b) FLAG-AT1R binding to yeast cells displaying nanobodies in Supplementary Data Fig. 1f, c) Biotinylated insect cell membrane polyspecificity reagent binding to yeast cells displaying nanobodies in Supplementary Data Fig. 1g. A final gate (P4) was included to eliminate extreme outliers.

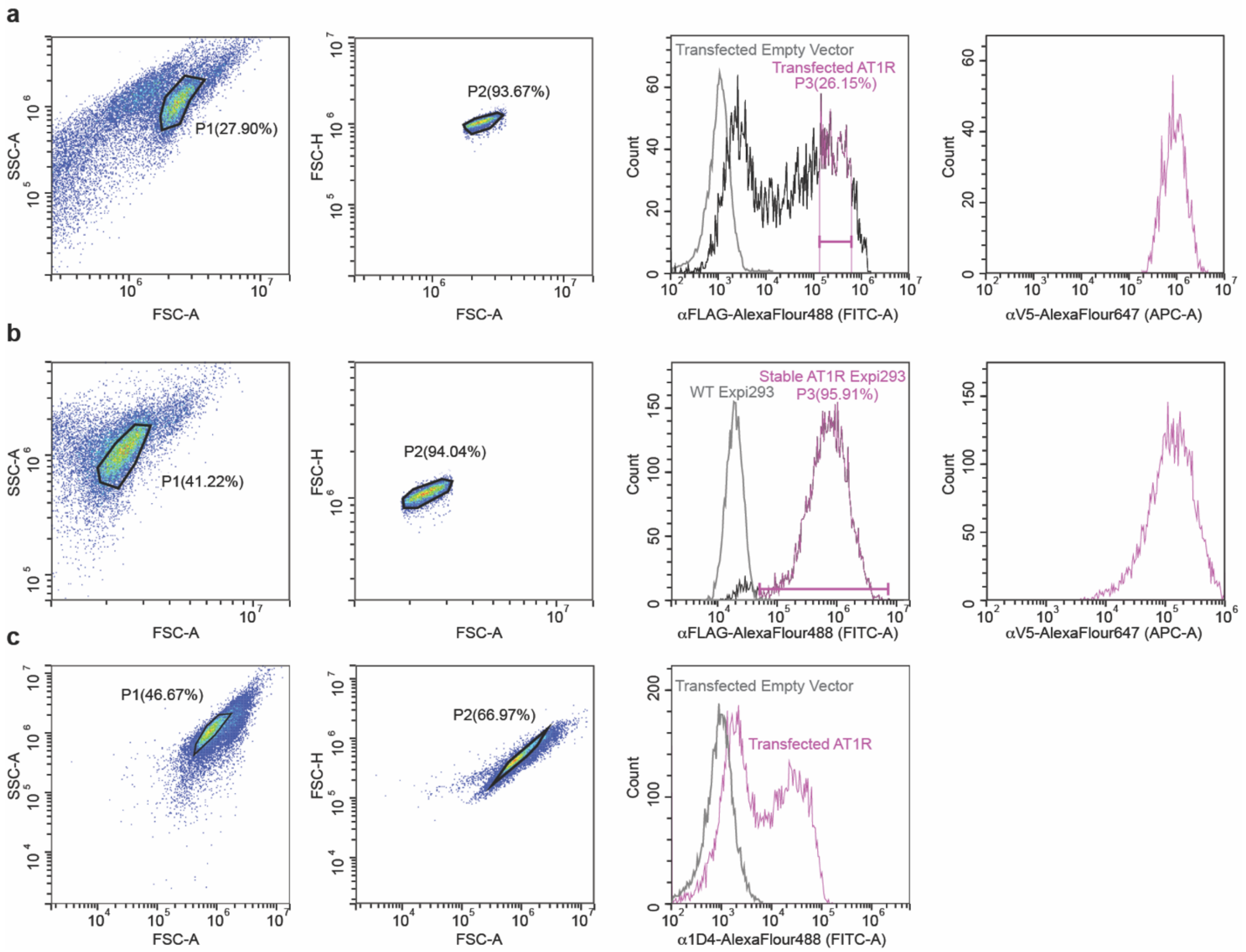

**Supplementary Data Figure 8.** Representative flow cytometry gating of mammalian cells. Flow cytometry plots illustrating gating strategies for a) Expi293 cells transiently transfected with FLAG-AT1R variants displayed in Fig. 3g, b) Expi293 cells stably expressing WT FLAG-AT1R displayed in Fig. 5f and Supplementary Data Fig. 5c. c) Fixed and permeabilized Expi293 cells transfects with C-terminally 1D4 tagged AT1R variants displayed in Supplementary Data Fig. 3a.

**Supplementary Data Table 1.** Nanobodies used in this study.

| Plasmid | Construct | Alternate Construct Name | Sequence |
|---|---|---|---|
| pMAS171 | AT118-A V5-His | AT118i4[13] | QVQLQESGGGLVQAGGSLRLSCAASGYIYRRYRMGWYRQAPGKEREFVAGINGGSSTNYADSVKGRFTISRDNAKNTVYLQMNSLKPEDTAVYYCAAYRIVWDLRVYWGQGTQVTVSSGKPIPNPLLGLDSTLEHHHHHH |
| pMAS177 | AT118-H V5-His | AT118i4h32[13,16] | EVQLVESGGGLVQPGGSLRLSCAASGYIYRRYRMGWYRQAPGKGREFVAAISGGSSTNYADSVKGRFTISRDNSKNTVYLQMNSLRAEDTAVYYCAAYRIVSDPRVYWGQGTQVTVSSGKPIPNPLLGLDSTLEHHHHHH |
| pMAS115 | AT118-L V5-His | | EVQLVESGGGLVQPGGSLRLSCAASGYIYSRYRMGWYRQAPGKGREFVAAISGGSSTNYADSVKGRFTISRDNSKNTVYLQMNSLRAEDTAVYYCAAYKIDSNPRVYWGQGTQVTVSSGKPIPNPLLGLDSTLEHHHHHH |
| pMAS515 | Anti-Fab nanobody FLAG-His | | QVQLQESGGGLVQPGGSLRLSCAASGRTISRYAMSWFRQAPGKEREFVAVARRSGDGAFYADSVQGRFTVSRDDAKNTVYLQMNSLKPEDTAVYYCAIDSDTFYSGSYDYWGQGTQVTVSSLEDYKDDKHHHHHH |
| pMAS517 | AT118-H MBP-His | | EVQLVESGGGLVQPGGSLRLSCAASGYIYRRYRMGWYRQAPGKGREFVAAISGGSSTNYADSVKGRFTISRDNSKNTVYLQMNSLRAEDTAVYYCAAYRIVSDPRVYWGQGTQVTVPPLVIWINGDKGYNGLAEVGKKFEKDTGIKVTVEHPDKLEEKFPQVAATGDGPDIIFWAHDRFGGYAQSGLLAEITPDKAFQDKLYPFTWDAVRYNGKLIAYPIAVEALSLIYNKDLLPNPPKTWEEIPALDKELKAKGKSALMFNLQEPYFTWPLIAADGGYAFKYENGKYDIKDVGVDNAGAKAGLTFLVDLIKNKHMNADTDYSIAEAAFNKGETAMTINGPWAWSNIDTSKVNYGVTVLPTFKGQPSKPFVGVLSAGINAASPNKELAKEFLENYLLTDEGLEAVNKDKPLGAVALKSYEEELAKDPRIAATMENAQKGEIMPNIPQMSAFWYAVRTAVINAASGRQTVDEALKDAQTLEHHHHHH |

| | | | |
|---|---|---|---|
| pMAS526 | Nb6 alpaca<br>ProteinC-His | | QRQLVESGGGLVQPGGSLRLSCAASG<br>TIFRLYDMGWFRQAPGKEREGVASITS<br>GGSTKYADSVKGRFIISRDNVKNTVYL<br>QMNSLEPEDTAVYYCNAEYRTGIWEE<br>LLDGWGKGTPVTVSSLEEDQVDPRLID<br>GKHHHHHH |
| pMAS430 | *IL2 signal sequence*-AT118-L – GSSG linker - Fc, **LALA** + **PG** | | *MYRMQLLSCIALSLALVTNS*EVQLVES<br>GGGLVQPGGSLRLSCAASGYIYSRYR<br>MGWYRQAPGKGREFVAAISGGSSTN<br>YADSVKGRFTISRDNSKNTVYLQMNSL<br>RAEDTAVYYCAAYKIDSNPRVYWGQG<br>TQVTVSSGGSGGGSGGGSGDKTHTC<br>PPCPAPE**AA**GGPSVFLFPPKPKDTLMI<br>SRTPEVTCVVVDVSHEDPEVKFNWYV<br>DGVEVHNAKTKPREEQYNSTYRVVSV<br>LTVLHQDWLNGKEYKCKVSNKAL**G**API<br>EKTISKAKGQPREPQVYTLPPSREEMT<br>KNQVSLTCLVKGFYPSDIAVEWESNG<br>QPENNYKTTPPVLDSDGSFFLYSKLTV<br>DKSRWQQGNVFSCSVMHEALHNHYT<br>QKSLSLSPGK |
| pMAS493 | *IL2 signal sequence*-AT118-L – GSSG linker -Fc, **LALA** + **PG**, I253**A**, H310**A**, H435**A**[1] | | *MYRMQLLSCIALSLALVTNS*EVQLVES<br>GGGLVQPGGSLRLSCAASGYIYSRYR<br>MGWYRQAPGKGREFVAAISGGSSTN<br>YADSVKGRFTISRDNSKNTVYLQMNSL<br>RAEDTAVYYCAAYKIDSNPRVYWGQG<br>TQVTVSSGGSGGGSGGGSGDKTHTC<br>PPCPAPE**AA**GGPSVFLFPPKPKDTLM<br>**A**SRTPEVTCVVVDVSHEDPEVKFNWY<br>VDGVEVHNAKTKPREEQYNSTYRVVS<br>VLTVL**A**QDWLNGKEYKCKVSNKAL**G**A<br>PIEKTISKAKGQPREPQVYTLPPSREE<br>MTKNQVSLTCLVKGFYPSDIAVEWES<br>NGQPENNYKTTPPVLDSDGSFFLYSKL<br>TVDKSRWQQGNVFSCSVMHEALHN**A**<br>YTQKSLSLSPGK |
| pMAS512 | *IL2 signal sequence*-AT118-L F47**T**, Y98**F** – GSSG linker -Fc **LALA** + **PG** | | *MYRMQLLSCIALSLALVTNS*EVQLVES<br>GGGLVQPGGSLRLSCAASGYIYSRYR<br>MGWYRQAPGKGRE**T**VAAISGGSSTN<br>YADSVKGRFTISRDNSKNTVYLQMNSL<br>RAEDTAVYYCAA**F**KIDSNPRVYWGQG<br>TQVTVSSGGSGGGSGGGSGDKTHTC<br>PPCPAPE**AA**GGPSVFLFPPKPKDTLMI<br>SRTPEVTCVVVDVSHEDPEVKFNWYV<br>DGVEVHNAKTKPREEQYNSTYRVVSV<br>LTVLHQDWLNGKEYKCKVSNKAL**G**API |

| | | | |
|---|---|---|---|
| | | | EKTISKAKGQPREPQVYTLPPSREEMT KNQVSLTCLVKGFYPSDIAVEWESNG QPENNYKTTPPVLDSDGSFFLYSKLTV DKSRWQQGNVFSCSVMHEALHNHYT QKSLSLSPGK |
| pMAS513 | *IL2 signal sequence*- AT118-L F47**T**, Y98**F** – GSSG linker -Fc, LAL**A** + P**G**, I253**A**, H310**A**, H435**A** [1] | | *MYRMQLLSCIALSLALVTNS*EVQLVES GGGLVQPGGSLRLSCAASGYIYSRYR MGWYRQAPGKGRE**T**VAAISGGSSTN YADSVKGRFTISRDNSKNTVYLQMNSL RAEDTAVYYCAA**F**KIDSNPRVYWGQG TQVTVSSGGSGGGSGGGSGDKTHTC PPCPAPE**AA**GGPSVFLFPPKPKDTLM **A**SRTPEVTCVVVDVSHEDPEVKFNWY VDGVEVHNAKTKPREEQYNSTYRVVS VLTVL**A**QDWLNGKEYKCKVSNKAL**G**A PIEKTISKAKGQPREPQVYTLPPSREE MTKNQVSLTCLVKGFYPSDIAVEWES NGQPENNYKTTPPVLDSDGSFFLYSKL TVDKSRWQQGNVFSCSVMHEALHN**A** YTQKSLSLSPGK |

[1]Fc Residue numbering is according to the Eu scheme[59].

**Supplementary Data Table 2.** Binding of AT118-H, AT118-L, and AT118-L-Fc fusion proteins to AT1R.

| Protein Construct | Kd (nM) |
|---|---|
| AT118-H (pMAS177) | 92.0 ± 6.0 |
| AT118-L (pMAS115) | 64.1 ± 7.6 |
| AT118-L Control Fc (pMAS430) | 29.0 ± 2.7 |
| AT118-L-Fc non-FcRn binding Fc (pMAS493) | 27.6 ± 2.0 |
| AT118-L F47T, Y98F Control Fc (pMAS512) | N.B. |
| AT118-L F47T, Y98F non-FcRn binding Fc (pMAS513) | N.B. |

The results are presented as mean Kd value ± S.E.M. from three independent experiments. N.B. means no binding over non-specific levels. pMAS# corresponds to construct described in Supplementary Table 1.

**Supplementary Data Table 3. Receptor construct sequences**

| Plasmid | Construct | Sequence |
|---|---|---|
| pMAS 85 | *Signal Sequence*-FLAG-AT1R | *MKTIIALSYIFCLVFA*DYKDDDDKILNSSTEDGIKRIQDDCPKAGRHNYIFVMIPTLYSIIFVVGIFGNSLVVIVIYFYMKLKTVASVFLLNLALADLCFLLTLPLWAVYTAMEYRWPFGNYLCKIASASVSFNLYASVFLLTCLSIDRYLAIVHPMKSRLRRTMLVAKVTCIIIWLLAGLASLPAIIHRNVFFIENTNITVCAFHYESQNSTLPIGLGLTKNILGFLFPFLIILTSYTLIWKALKKAYEIQKNKPRNDDIFKIIMAIVLFFFFSWIPHQIFTFLDVLIQLGIIRDCRIADIVDTAMPITICIAYFNNCLNPLFYGFLGKKFKRYFLQLLKYIPPKAKSHSNLSTKMSTLSYRPSDNVSSSTKKPAPCFEVE |
| pMAS 625 | *Signal Sequence*-AT1RΔ320-3C-Rho1D4 | *MKTIIALSYIFCLVFA*ILNSSTEDGIKRIQDDCPKAGRHNYIFVMIPTLYSIIFVVGIFGNSLVVIVIYFYMKLKTVASVFLLNLALADLCFLLTLPLWAVYTAMEYRWPFGNYLCKIASASVSFNLYASVFLLTCLSIDRYLAIVHPMKSRLRRTMLVAKVTCIIIWLLAGLASLPAIIHRNVFFIENTNITVCAFHYESQNSTLPIGLGLTKNILGFLFPFLIILTSYTLIWKALKKAYEIQKNKPRNDDIFKIIMAIVLFFFFSWIPHQIFTFLDVLIQLGIIRDCRIADIVDTAMPITICIAYFNNCLNPLFYGFLGKKFKRYFLQLLKYGGSSLEVLFQGPTETSQVAPA |
| pMAS 514 | *Signal Sequence*-AT118-H-proteinC-AT1R-226-BRIL-A2A/Frizzled5 H6-234-Δ320-3C-Rho1D4 | *MKTIIALSYIFCLVFA*EVQLVESGGGLVQPGGSLRLSCAASGYIYRRYRMGWYRQAPGKGREFVAAISGGSSTNYADSVKGRFTISRDNSKNTVYLQMNSLRAEDTAVYYCAAYRIVSDPRVYWGQGTQVTVSSLEGGSGEDQVDPRLIDGKILNSSTEDGIKRIQDDCPKAGRHNYIFVMIPTLYSIIFVVGIFGNSLVVIVIYFYMKLKTVASVFLLNLALADLCFLLTLPLWAVYTAMEYRWPFGNYLCKIASASVSFNLYASVFLLTCLSIDRYLAIVHPMKSRLRRTMLVAKVTCIIIWLLAGLASLPAIIHRNVFFIENTNITVCAFHYESQNSTLPIGLGLTKNILGFLFPFLIILTSYTLIWKALKKAYDLEDNWETLNDNLKVIEKADNAAQVKDALTKMRAAALDAQKATPPKLEDKSPDSPEMKDFRHGFDILVGQIDDALKLANEGKVKEAQAAAEQLKTTRNAYIQKYLERARSTLDKLNDDIFKIIMAIVLFFFFSWIPHQIFTFLDVLIQLGIIRDCRIADIVDTAMPITICIAYFNNCLNPLFYGFLGKKFKRYFLQLLKYGGSSLEVLFQGPTETSQVAPA |
| pMAS 516 | *Signal Sequence*-FLAG-AT1R226-BRIL-A2A/Frizzled5 H6-234-Δ320 | *MKTIIALSYIFCLVFA*DYKDDDDKILNSSTEDGIKRIQDDCPKAGRHNYIFVMIPTLYSIIFVVGIFGNSLVVIVIYFYMKLKTVASVFLLNLALADLCFLLTLPLWAVYTAMEYRWPFGNYLCKIASASVSFNLYASVFLLTCLSIDRYLAIVHPMKSRLRRTMLVAKVTCIIIWLLAGLASLPAIIHRNVFFIENTNITVCAFHYESQNSTLPIGLGLTKNILGFLFPFLIILTSYTLIWKALKKAYDLEDNWETLNDNLKVIEKADNAAQVKDALTKMRAAALDAQKATPPKLEDKSPDSPEMKDFRHGFDILVGQIDDALKLANEGKVKEAQAAAEQLKTTRNAYIQKYLERARSTLDKLNDDIFKII |

| | | |
|---|---|---|
| | | MAIVLFFFFSWIPHQIFTFLDVLIQLGIIRDCRIADIVDTAMPITICIAYFNNCLNPLFYGFLGKKFKRYFLQLLKY |
| pMAS 480 | *Signal Sequence*-AT118-H-proteinC-AT1R-217-Kappa Opioid ICL3-244-Δ320-3C-Rho1D4 | *MKTIIALSYIFCLVFA*EVQLVESGGGLVQPGGSLRLSCAASGYIYRRYRMGWYRQAPGKGREFVAAISGGSSTNYADSVKGRFTISRDNSKNTVYLQMNSLRAEDTAVYYCAAYRIVSDPRVYWGQGTQVTVSSLEGGSGEDQVDPRLIDGKILNSSTEDGIKRIQDDCPKAGRHNYIFVMIPTLYSIIFVVGIFGNSLVVIVIYFYMKLKTVASVFLLNLALADLCFLLTLPLWAVYTAMEYRWPFGNYLCKIASASVSFNLYASVFLLTCLSIDRYLAIVHPMKSRLRRTMLVAKVTCIIIWLLAGLASLPAIIHRNVFFIENTNITVCAFHYESQNSTLPIGLGLTKNILGFLFPFLIILTSYTLMILRLKSVRLLSGSREKDRNLRRITRLVLAIVLFFFFSWIPHQIFTFLDVLIQLGIIRDCRIADIVDTAMPITICIAYFNNCLNPLFYGFLGKKFKRYFLQLLKYGGSSLEVLFQGPTETSQVAPA |

### Supplementary Data Table 4. Cryo-EM Data Collection

### Cryo-EM data collection, refinement and validation statistics

| | AT118-H AT1R<br><br>(EMDB-41248)<br>(PDB 8TH3) | AT118-L AT1R<br>Losartan<br>(EMDB-41249)<br>(PDB 8TH4) |
|---|---|---|
| **Data collection and processing** | | |
| Magnification | 105,000 | 105,000 |
| Voltage (kV) | 300 | 300 |
| Electron exposure (e–/Å$^2$) | 64.2 | 64.1 |
| Defocus range (μm) | -0.8 to -1.8 | -0.8 to -1.8 |
| Pixel size (Å) | 0.83 | 0.83 |
| Symmetry imposed | C1 | C1 |
| Initial particle images (no.) | 6,309,488 | 518,841 |
| Final particle images (no.) | 478,571 | 238,621 |
| Map resolution (Å) | 2.9 | 3.25 |
| FSC threshold | 0.143 | 0.143 |
| **Refinement** | | |
| Initial model used (PDB code) | 4ZUD, 7T83, 6WW2 | 4ZUD, 7T83, 6WW2 |
| Model resolution (Å) | 2.8, 2.1, 3.7 | 2.8, 2.1, 3.7 |
| Map sharpening *B* factor (Å$^2$) | 123 | 105 |
| Model composition | | |
| Non-hydrogen atoms | 5770 | 5702 |
| Protein residues | 720 | 722 |
| Ligands | 2 | 1 |
| R.m.s. deviations | | |
| Bond lengths (Å) | 0.003 | 0.003 |
| Bond angles (°) | 0.53 | 0.55 |
| Validation | | |
| MolProbity score | 2.26 | 1.84 |
| Clashscore | 8.6 | 9.2 |
| Poor rotamers (%) | 3.6 | 0.2 |
| Ramachandran plot | | |
| Favored (%) | 94.5 | 94.9 |
| Allowed (%) | 5.5 | 5.1 |
| Disallowed (%) | 0 | 0 |

**Supplementary Data Table 5. Olmesartan Derivative Compound Validation**

| VendorID | Compound | Calculated m/z [M+H] | Found m/z [M+H] | Purity (%)[1] |
|---|---|---|---|---|
| SL-3662 | Olmesartan Methyl Ester | 461.2296 | 461.2294 | 98.6 |
| SL-3663 | Dehydro Olmesartan | 429.2034 | 429.2036 | 97.9 |
| SL-3665 | Olmesartan Methyl Ketone | 445.2347 | 445.2346 | 98.3 |

[1] % purity based on HPLC

**Supplementary Data Table 6. Primers used in this study**

| | |
|---|---|
| P94 | GAGGTGCAGCTGGTAGAAAGCGGCGGCGGCCTGGTGCAGCCAGGCGGCAGC |
| P95 | GCCGCTCGCCGCGCAGCTCAGGCGCAGGCTGCCGCCTGGCTGC |
| P96 | GCGGCGAGCGGCDMCDWTDACARSSRAKMTASAATGGGCTGGTATCGCCAGG |
| P97 | TTCGCGACCTTTGCCCGGCGCCTGGCGATACCAGCCCAT |
| P98 | CCGGGCAAAGGTCGCGAATTCGTTGCCGCTATTTCAGGCGGTYCATCCACAAACTAT GCGGATAGCGTGAAAGGCC |
| P99 | GTTTTTCGAGTTATCGCGGCTAATGGTAAAGCGGCCTTTCACGCTATCCGCATA |
| P100 | AGCCGCGATAACTCGAAAAACACCGTGTATCTGCAGATGAACAGCCTGCGTGCC |
| P101 | CGCGCAATAATACACCGCGGTATCTTCGGCACGCAGGCTGTTCATCTGCAGA |
| P102 | CCGCGGTGTATTATTGCGCGGYCDATAVGANCGHCRRTRRCCCGASAGWCTATTGG GGCCAGGGCACC |
| P103 | GCTGCTCACGGTCACCTGGGTGCCCTGGCCCCAATA |
| P104 | GAAGGTGTTCAATTGGACAAGAGAGAAGCTGACGCAGAGGTGCAGCTGGTAGAAAG CGGC |
| pYDS Rev1 | TACTGATGCTTCTGTAGAGGGTGAGGATGTTTGAGCGTAATCTGGAACATCGTATG GGTAGGATCCGCTGCTCACGGTCACCTGGGTGCC |
| pYDS Fwd2 | ACCACCATCGCTTCTATCGCTGCTAAGGAAGAAGGTGTTCAATTGGACAAGAGAG AAGC |
| pYDS Rev2 | TACTGATGCTTCTGTAGAGGGTGAGG |
| pYDS Fwd3 | ACCACCATCGCTTCTATCGCTGCTAAG |
| P209 | GTTCAATTGGACAAGAGAGAAGCT |
| P210 | GTAATCTGGAACATCGTATGGGTA |
